# Influence of DNA extraction methods on microbiome and resistome analysis in activated sludge

**DOI:** 10.1101/2023.06.26.546617

**Authors:** David Calderón-Franco, Dana Kok, Renzo Dukker, Ben Abbas, Joana Abreu-Silva, Jaqueline Rocha, Marco Antonio Lopez Marin, Rebeca Pallarés Vega, Stanislav Gajdos, Mythili Ananth, Seeram Apoorva, Ivan Karpisek, Milada Solcova, Lucia Hernández Leal, Jan Bartacek, Célia Manaia, Sabina Purkrtova, David G. Weissbrodt

## Abstract

Amplicon sequencing, metagenomics, and quantitative polymerase chain reaction (qPCR) are commonly used techniques to analyse microorganisms and antibiotic resistance genes (ARGs) in activated sludge from wastewater treatment plants (WWTPs). However, the lack of workflow harmonisation poses challenges in comparing measurements across studies and research groups. To address this issue, we examined the impact of DNA extraction procedures on 16S rRNA gene amplicon sequencing, shotgun metagenomics, and qPCR analyses of activated sludge by combining two widely used DNA extraction kits (PowerSoil and FastDNA) and two commonly employed disruption instruments (bead-beater and vortex) through a 2×2 factorial experimental design involving four groups of three analysts performing DNA extractions in triplicates. Our findings revealed significant differences in DNA yield, purity, and reproducibility of amplicon sequencing profiles among the extraction kits. Operator variability also influenced the results. We compared microbiome profiles obtained by amplicon sequencing and metagenomics and observed that bead-beating introduced more variability among triplicates compared to vortexing. The combinations of extraction kits and disruption instruments impacted the relative abundances of specific phyla such as *Actinobacteriota*, *Bacteroidota*, and *Nitrospirota*. For resistome analysis, we employed metagenomics for high-resolution profiling and qPCR for high-sensitivity detection of ARGs. The compositions and diversities of resistome datasets were not significantly affected by the choice of extraction kits and disruption instruments. Although using the same method is ideal for accurate comparisons, our results suggest that acceptable reproducibility can still be achieved when using different methods. This finding encourages the implementation of ARG monitoring in wastewater treatment processes. However, it is important to consider biases introduced by DNA extraction workflows when designing analytical studies, interpreting their results, and comparing their findings. Striving for more harmonised molecular workflows is crucial in the field of wastewater microbiology and engineering.

Graphical abstract

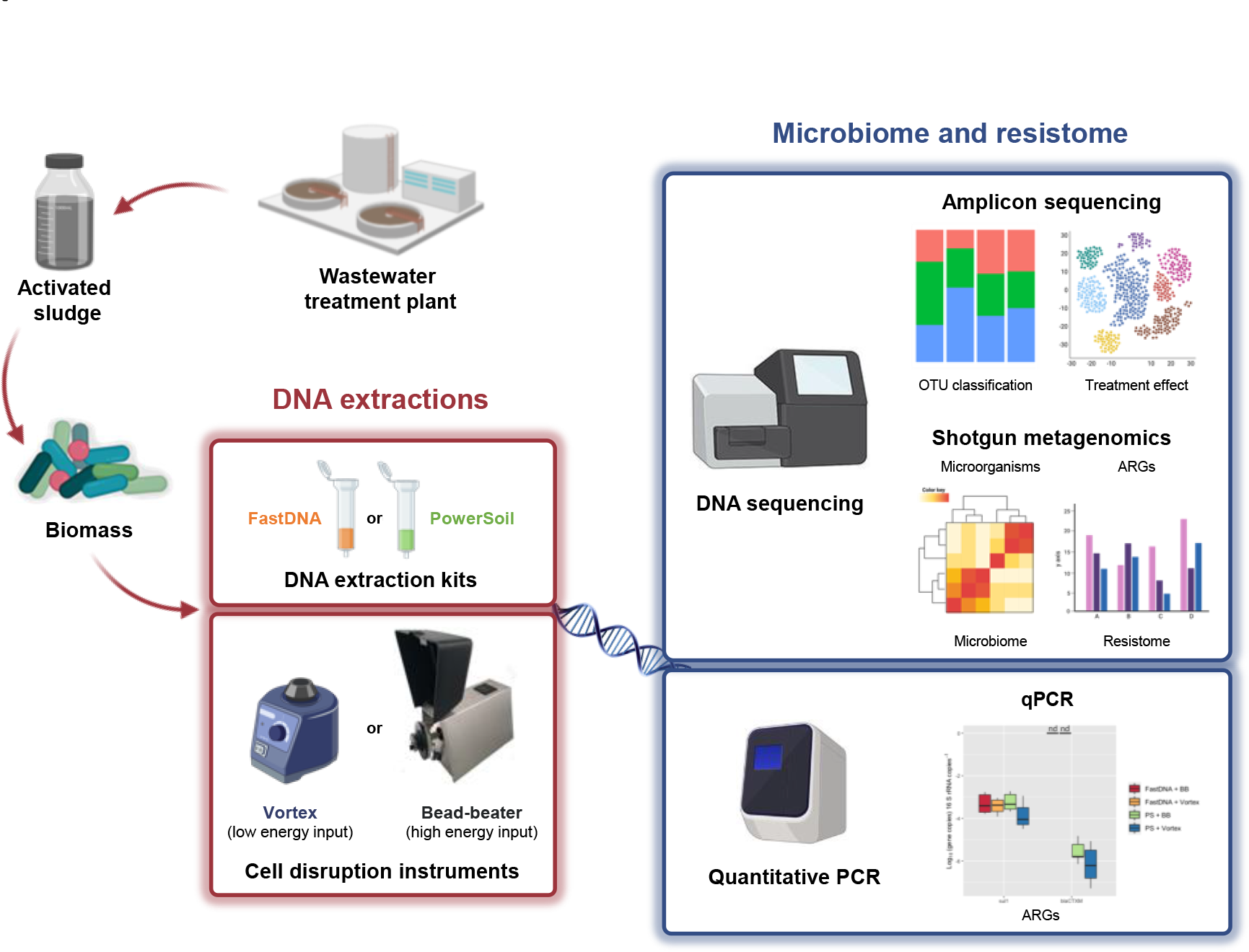

## Introduction

Antimicrobial resistance (AMR) has become a major concern, posing a significant threat to the effectiveness of antibiotics in both the medical field and the food industry. Since their discovery in the early 20th century, antibiotics have played a crucial role in treating infectious diseases. However, the emergence and spread of AMR have undermined the efficacy of these life-saving drugs.

To mitigate environmental pollution and address the growing demand for clean water, sewer systems and wastewater treatment plants (WWTPs) are essential (Anthony et al., 2020). These facilities are designed to remove pathogens and nutrients from wastewater, ensuring the safety of water resources. In the context of the One Water – One Health approach, new challenges arise from the diverse forms of pollution within wastewater catchment areas, influenced by regional socioeconomic factors and cultural practices. Constantly evolving analytical methods are employed to identify and understand these pollution sources (Miłobedzka et al., 2022).

In addition to micropollutants, nanoparticles, and microplastics, the presence of antibiotic resistance genes (ARGs) and antibiotic resistant bacteria (ARB) has become a major concern for water authorities (Ju et al., 2016; Mintenig et al., 2017; Hendriksen et al., 2019). Wastewater and wastewater treatment plants (WWTPs), as integral components of the urban water cycle, are recognised as reservoirs for ARB and ARGs (Rizzo et al., 2013; Anthony et al., 2020) and serve as interfaces for their dissemination into the environment (Liu et al., 2019). Consequently, there is an increasing focus on studying the emission, amplification, and removal of antibiotic resistance determinants within urban water systems.

The identification and quantification of ARB and ARGs in wastewater environments are essential tasks for surveillance programs (Manaia et al., 2018; Pallarés-Vega et al., 2020). However, the lack of harmonisation and standardisation in molecular biology workflows presents challenges when comparing studies conducted by different research groups (Rocha et al., 2020).

Commercial DNA extraction kits commonly used for other environmental sample types, such as soil, water, or generic biofilms, are often applied to extract DNA from activated sludge. However, activated sludge is a complex and diverse matrix consisting of various microbial populations with different cellular compositions and aggregation states, presenting a challenge for extraction methods to obtain a representative DNA pool (McIlroy et al., 2009; Guo and Zhang, 2013). DNA extraction methods play a crucial role not only in the detection and quantification of genes by quantitative PCR (qPCR) (Rocha et al., 2020; Agrawal et al., 2021) but also in the profiling of microbial communities such as by 16S rRNA gene amplicon sequencing (Albertsen et al., 2015), influencing the results obtained. It is essential to identify and consider biases at the lowest taxonomic level possible, as ecologically relevant traits often exhibit phylogenetic conservation (Martiny et al., 2015; Dueholm et al., 2020).

DNA sequencing techniques, such as amplicon sequencing and shotgun metagenomics, have revolutionised the profiling of microbiome and resistome in wastewater systems, offering cost-efficient and culture-independent approaches (Schmieder and Edwards, 2012; Li et al., 2013; Wang et al., 2013; Guo et al., 2017; Cerruti et al., 2021; Calderón-Franco et al., 2022). These methods provide high-resolution insights into population and genetic profiles, complementing traditional qPCR workflows commonly used for measuring selected genes. However, qPCR measurements for ARGs are susceptible to primer bias and limited coverage of ARG targets, whereas sequencing outputs are influenced by library preparation (Schmieder and Edwards, 2012; Mullany, 2014; Li et al., 2018; Zhou et al., 2019; Albertsen et al., 2015). Additionally, both amplicon-based and sequencing results can be affected by DNA extraction methods, which pose significant challenges to the reliability and comparability of molecular findings (Guo and Zhang, 2013; Albertsen et al., 2015; Rocha et al., 2019; Gołębiewski and Tretyn, 2020; Agrawal et al., 2021).

16S rRNA gene amplicon sequencing is commonly used to analyse microbial communities in environmental studies. However, its ability to classify taxa beyond the genus level is limited due to the short length of the sequenced amplicons (Janda and Abbott, 2007; Kim and Chun, 2014; Weissbrodt et al., 2014; Albertsen et al., 2015; Dueholm et al., 2020). Additional challenges arise from various factors such as closely related gene sequences, variations in the number of gene copies across microbial genomes, biases introduced by PCR amplification, nomenclature issues resulting from multiple genomovars assigned to single species, establishing new taxa, or inadequate reference databases (Case et al., 2007). Another limitation is its inability to differentiate between DNA from live and dead cells (Li et al., 2017), whereas treatments with ethidium monoazide bromide or propidium monoazide can selectively target DNA from dead cells and prevent its amplification in PCR (Siefring et al., 2008; Vondrakova et al., 2018).

Despite the importance of extracting informational macromolecules, such as nucleic acids, proteins, lipids, and polysaccharides, from cells, limited research has been conducted on the effect of different DNA extraction workflows on both the microbiome and resistome of activated sludge (Bushon et al., 2010; Vanysacker et al., 2010; Guo and Zhang, 2013; Albertsen et al., 2015; Li et al., 2018). Therefore, it is necessary to address these limitations and harmonise molecular workflows to enable reliable comparisons of results across different laboratories.

Within the training framework of the European Twinning project REPARES (http://repares.vscht.cz/), our study focused on investigating the effects of various DNA extraction methods, involving different kits and disruption instruments, on the molecular analysis of microorganisms and ARGs in activated sludge. We employed 16S rRNA gene amplicon sequencing (for bacterial community profiling), metagenomics (for microbiome and resistome analysis), and qPCR (for targeted gene analysis). Our ultimate goal was to contribute to the international efforts aimed at establishing harmonised molecular protocols for profiling the microbiome and resistome in wastewater environments.

## Material and Methods

### Sample collection from the activated sludge tank of a WWTP

Biological samples were collected from the urban WWTP Harnaschpolder (Waterboard Delftland, The Netherlands) operated for full biological nutrient removal. A volume of 1 L of mixed liquor was collected as a grab sample from the activated sludge tank under dry weather flow (i.e., no recent rainfall and variations in the hydraulic retention time). The raw activated sludge sample was stored at 4 °C in a time frame of less than 4 h prior to DNA extractions. The activated sludge comprised 4.5 g of total, 1.5 g of inorganic (27%) and 3.3 g of volatile suspended solids (73%) per litre.

### DNA extraction, quantification, and quality

DNA extraction from the activated sludge sample was performed using a 2×2 factorial experimental design combining 2 DNA extraction kits and 2 disruption instruments, by 4 groups of 3 analysts each, enabling to also address operator effects (**Table 1**).

**Table 1.** The 2×2 factorial experimental design combining 2 DNA extraction kits and 2 disruption instruments was performed by 4 groups of 3 analysts. Each combination was conducted on biological triplicates, resulting in a total of 12 DNA extracts analysed by amplicon sequencing, metagenomics, and qPCR. Acronyms used in figures for kits and instruments are given in brackets.

| Group | Extraction kit | Disruption instrument |
| --- | --- | --- |
| 1 | FastDNA (FD) | Bead-beater (BB) |
| 2 | FastDNA (FD) | Vortex (VT) |
| 3 | PowerSoil (PS) | Bead-beater (BB) |
| 4 | PowerSoil (PS) | Vortex (VT) |

The widely used FastDNA® Spin kit for soil (MP Biomedicals, USA) and PowerSoil® DNA isolation kit (MoBio, USA; now produced by Qiagen, USA) were used according to manufacturers’ protocols. An amount of 0.250 g of wet biomass was collected after centrifugation of the activated sludge samples (4.5 g TSS L^-1^) at 13000 × g during 2 min and resuspended in the respective lysis or bead solutions before extraction.

DNA extraction with each kit was tested using two widely used instruments that disrupt cells with high (*i.e.*, bead-beater) and low (*i.e.*, vortex) energy. Bead-beating was performed on a Mini-Beadbeater-24 (BioSpec Products, USA) at default intensity settings following general operation recommendations: 10 minutes for the PowerSoil® DNA isolation kit and 40 seconds for the FastDNA® isolation kit. Vortex was performed on a Vortex-Genie 2 (MO BIO Laboratories, USA) at maximum intensity, following the same operation recommendations as for bead-beating: 10 minutes for the PowerSoil® DNA isolation kit and 40 seconds for the FastDNA® isolation kit. Each combination of DNA extraction conditions was performed in triplicate.

The DNA yield, purity, and integrity were measured on the triplicated DNA extracts. The purity of the DNA extracts was evaluated with a Nanodrop® spectrophotometer using the ratios of absorbance measurements at 230, 260 and 280 nm (*i.e.*, A_230_/A_260_ and A_260_/A_280_) (Thermo Fisher Scientific, USA). The integrity (DNA fragment size) of extracted DNA was evaluated using a 1% (w/v) agarose gel electrophoresis (Sigma-Aldrich, United Kingdom) in 1⊆ tris-acetate-EDTA (TAE) buffer. The DNA concentration was measured fluorometrically with Qubit R dsDNA assays (Thermo Fisher Scientific, USA).

### 16S rRNA gene amplicon sequencing

#### Preparation and sequencing of amplicon libraries

All 12 DNA extracts were sent on dry ice to Novogene Ltd. (Hongkong, China) for paired-end amplicon sequencing of the V3-V4 region of the 16S rRNA gene pool, using the standard protocol of the sequencing facility. Barcoded forward 341f (5’-CCTAYGGGRBGCASCAG-3’) and reverse 806r (5’-GGACTACNNGGGTATCTAAT-3’) oligonucleotide primers were used for PCR to produce amplicons of 470 bp (*i.e.*, *Escherichia coli* position 341-806).

The PCRs were prepared by the sequencing facility according to its internal protocol. In short, the concentration and purity of the DNA extracts were evaluated on 1% agarose gels. The DNA extracts were diluted to 1 ng μL^-1^ using DNase/RNase-free water. The PCRs were conducted in volumes of 30 μL comprising a Phusion® High-Fidelity PCR Master Mix (New England Biolabs) (15 μL), forward and reverse primers (0.2 μmol L^-1^ each), template DNA (10 ng), and DNase/RNase-free water (completed to total reaction volume). The PCR thermal cycling program consisted of an initial denaturation (1 minute at 98°C) followed by 30 cycles of denaturation (10 seconds at 98°C, annealing (30 seconds at 50°C) and elongation (60 seconds at 72°C), and was terminated by a final elongation (5 minutes at 72°C).

After quality control by agarose gel electrophoresis and quantification, the PCR products were mixed in equidensity ratios. The pool of mixed PCR products was purified with a GeneJET Gel Extraction Kit (Thermo Fisher Scientific, USA). The sequencing library was generated using a NEBNext Ultra DNA Library Prep Kit for Illumina (New England BioLabs, USA). The concentration and quality of the library was analysed with a Qubit 2.0 Fluorometer (Thermo Fisher Scientific, USA) and a 2100 Bioanalyzer System (Agilent Technologies, USA).

The V3-V4 16S rRNA gene amplicon sequencing library was sequenced on a MiSeq benchtop platform (Illumina, USA) and datasets of 250 bp paired-end reads were generated. The amplicon sequencing datasets will be made available on NCBI.

#### Bioinformatics processing of amplicon sequences: paired-end reads assembly and quality control

Paired-end reads were divided by sample based on their unique barcode. Primer and barcode sequences were truncated from all reads. To acquire longer reads, FLASH was used to merge the paired-end reads. Raw sequences were cleaned using the QIIME pipeline (Bolyen et al., 2019). To remove chimera sequences, the cleaned sequences were compared with the Genomes OnLine Database (GOLD) v.8 (Mukherjee et al., 2021) through the UCHIME algorithm (Edgar et al., 2011). After the removal of chimeric sequences, the resulting sequences were used for taxonomic assignment. Operational taxonomic units (OTUs) were generated using the Uparse software (Edgar, 2013). The threshold to cluster sequences to the same OTU was set at 97%. To assign taxonomic rank to representative sequences of OTUs, the RDP classifier (naive Bayesian classifier) (Wang et al., 2007) and ARB-SILVA database (Quast et al., 2013) were used.

#### Numerical ecology analyses of amplicons sequencing datasets

OTU diversity, multivariate numerical analyses, and statistical analyses were performed with the R packages Phyloseq (McMurdie et al., 2013) and Ampvis 2 (Andersen et al., 2018). All plots were produced with the ggplot2 R package (Wickham, 2009). The Phyloseq package was used to calculate the alpha diversity indices: number of observed OTUs, Shannon, and Simpson of each sample.

To visualise relative abundance values in sample groups, heatmaps from different phylogenetic levels were produced using Ampvis 2 (Andersen et al., 2018). Phyloseq was used to make bar-plots of relative abundances of each sample at different phylogenetic levels. To allow comparison between samples with different sequencing depths, the median sequencing depth normalisation was used.

Non-metric dimensional scaling (NMDS) and principal coordinates analysis (PCoA) were used to visualise the differences in the observed microbial community compositions between samples (Ampvis 2 R package). NMDS and PCoA were performed using the Bray-Curtis dissimilarity measure. A principal component analysis (PCA) was used to visualise the consistency of principal components and numerical differences of major taxa between sample groups (Ampvis 2 R package).

### Metagenomic analysis of microbiome and resistome compositions

The compositions of the microbiomes and resistomes of the 12 total DNA extracts obtained from the activated sludge sample were analysed at high resolution following the metagenomics method that we used in earlier studies (Calderón-Franco et al., 2022; Calderón-Franco et al., 2023). The method is summarised hereafter.

#### Preparation and sequencing of metagenomics libraries

The same 12 DNA extracts sent to Novogene Ltd. (China) were used to prepare and sequence metagenome libraries, according to the internal protocol of the facility.

An amount of 1 μg of DNA was used per extract as input material. The NEBNext Ultra DNA Library Prep Kit for Illumina (New England BioLabs, USA) was used to prepare the metagenomic libraries. Index codes were included to attribute sequences to each sample. The DNA templates were fragmented to 350 bp by sonication followed by end-polished, A-tailed, and ligated with the full-length adaptor for Illumina sequencing with further PCR to insert the sequence adapters. The PCR products were purified on AMPure XP magnetic beads (Beckman Coulter, USA).

The tagged libraries were pooled and sequenced and sequenced on a HiSeq PE150 system (Illumina, USA), producing metagenomics datasets of 26.5±0.5 millions of 150 bp paired-end reads per sample. The characteristics of the sequencing datasets are given in **Table S1** in the supplementary information. The metagenomics datasets will be made available via NCBI.

#### Microbiome analysis

The quality of the acquired Illumina reads was assessed by FastQC (v.0.11.9) with default parameters (Andrews, 2010) and visualised with MultiQC (v.1.0) (Ewels, 2016). Low-quality paired-end reads were trimmed and filtered using Trimmomatic (v.0.39) on paired-end mode (Bolger et al., 2014). The clean sequence reads were classified to microbial taxonomies using Kraken2 (v.2.0) (Wood et al., 2019) with default parameters and its underlying database of all complete bacterial, archaeal, and viral genomes retrieved from the NCBI Refseq database (O’Leary et al., 2016). Kraken2 divided the reads into k-mers and matched them with the NCBI database. The absolute abundances of taxa were computed as the number of k-mers aligned to a specific phylotype, and their relative abundances by normalising to the total number of k-mers aligned in each sample. Absolute and relative abundances were calculated using the package Pavian (v.1.2.0) (Breitwieser and Salzberg, 2020) in R (R Core Team, 2020).

A non-metric multidimensional scaling (NMDS) ordination was computed in R to reduce the dimensions and visualise in two dimensions the distances between the metagenomics profiles of the 12 DNA extracts resulting from the 4 different DNA extraction combinations performed in triplicates. NMDS was applied to the taxonomic data at phylum and genus levels. Graphs were made with RStudio (version 1.3.1093).

#### Resistome analysis

To annotate ARGs, the clean sequence reads obtained from the metagenomic datasets were aligned by the BLASTp tool to the Comprehensive Antibiotic Resistance Database (CARD) (Alcock et al., 2020) with a cut-off e-value <1·10^-5^ and sequence identity >90%. The relative abundances of the identified sequences of ARG (sub)types in a metagenome were expressed as parts per million (ppm, 1 in 10^6^ sequences) of the total amounts of ARG-like sequences detected.

The Shannon index for the absolute amount of ARGs in the resistome was calculated for each different DNA extraction combination using the Vegan package (Oksanen et al., 2019) in R (R Core Team, 2020). To determine how both kits and extractions influence the detected amount of ARGs, a multiple linear regression was fit in which the Shannon index represents the dependent variable, and the kit and disruption instrument both explanatory variables.

### Quantitative polymerase chain reaction (qPCR) analysis of selected ARGs

qPCR was conducted on the two ARGs *sul1* and *bla*_CTX-M_ in addition to the bacterial 16S rRNA gene (**Table 2**) from all DNA extracts.

**Table 2.**
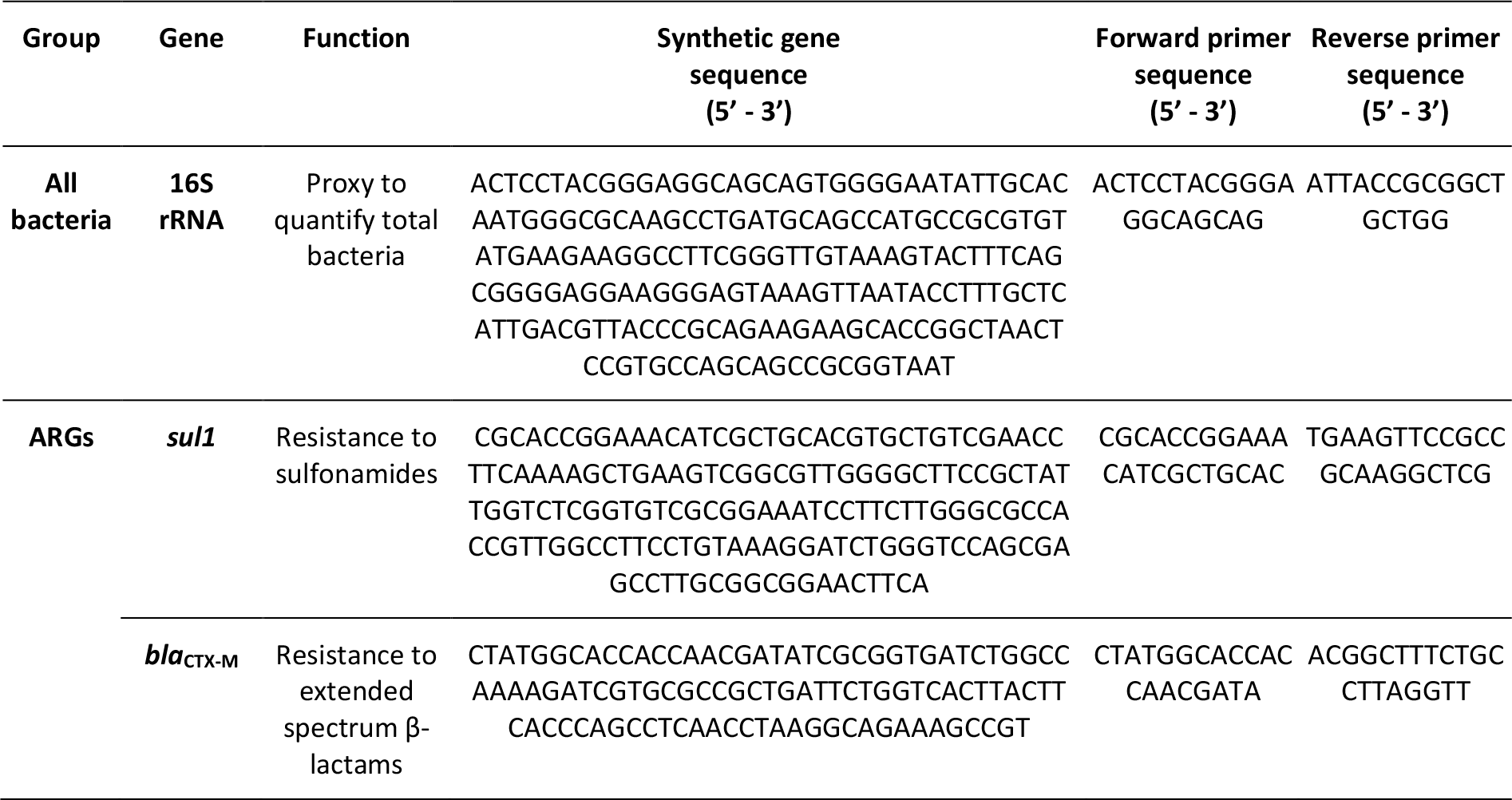
Genes analysed by qPCR from the 12 DNA extracts obtained from the activated sludge sample. Genes targeted two groups of interest, namely all bacteria (16S rRNA gene) and the two antibiotic resistance genes (ARGs) *sul1* and *bla*_CTX-M_. Synthetic partial DNA sequences of the 16S rRNA gene and of the two ARGs were retrieved from ResFinder to generate the standard curves for qPCR. The 5’-3’ sequences are indicated for the sets of oligonucleotide primers used for qPCR.

qPCR reactions were conducted in a total volume of 20 µL, including IQTM SYBR green supermix 1⊆ (Bio-Rad, USA), using the forward and reverse primers summarised in **Table 2**. A total of 2 µL of DNA template was added to each reaction, and the reaction volume was completed to 20 µL with DNase/RNase free water (Sigma Aldrich, UK). All reactions were performed in a qTOWER3 Real-time PCR machine (Westburg, DE) according to the following cycling conditions: 95°C for 5 minutes followed by 40 cycles at 95°C for 15 seconds and 60°C for 30 seconds. The annealing temperatures were 60°C for both the 16S rRNA and *bla*_CTX-M_ genes, and 65°C for *sul1*.

To check the specificity of the reaction, a melting curve of each amplicon was performed from 65 to 95°C at a temperature gradient of +0.5 °C (5 s)^-1^. Synthetic DNA fragments (IDT, USA) containing each of the target genes were used as a positive control to create the standard curves (**Table 2**). Serial 10-fold dilutions of gene fragments were performed in sheared salmon sperm DNA 5 µg mL^-1^ (m/v) (Thermofisher, LT) diluted in Tris-EDTA (TE) buffer at pH 8.0. Every DNA extract was analysed in technical triplicates. Standard curves were included in each qPCR plate with at least 6 serial dilutions points and each in technical duplicates. An average standard curve was created for each gene set based on the standard curves from every run. Gene concentration values were then calculated using the standard curve.

The effects of the different DNA extraction treatments on qPCR results were addressed by an analysis of variance (ANOVA) provided in **Table S2** in the supplementary information.

## Results

### Effect of commercial extraction kits and disruption instruments on DNA yield and quality

The impact of 2×2 combinations of commercial extraction kits and disruption instruments on DNA yield and quality was evaluated during the DNA extraction process from fresh activated sludge. The evaluation was conducted by four groups, each comprising three analysts.

DNA extraction performed using the PowerSoil kit resulted in significantly higher concentrations (129-236 ng µL^-1^) compared to the FastDNA kit (8-27 ng µL^-1^) (**Table 3**). This observation was consistent across both disruption instruments employed, bead-beater or vortex. The DNA purity was not compromised by neither the extraction kits nor the disruption instruments: the A_260_/A_280_ absorbance ratio ranged between 1.7-1.9.

**Table 3.**
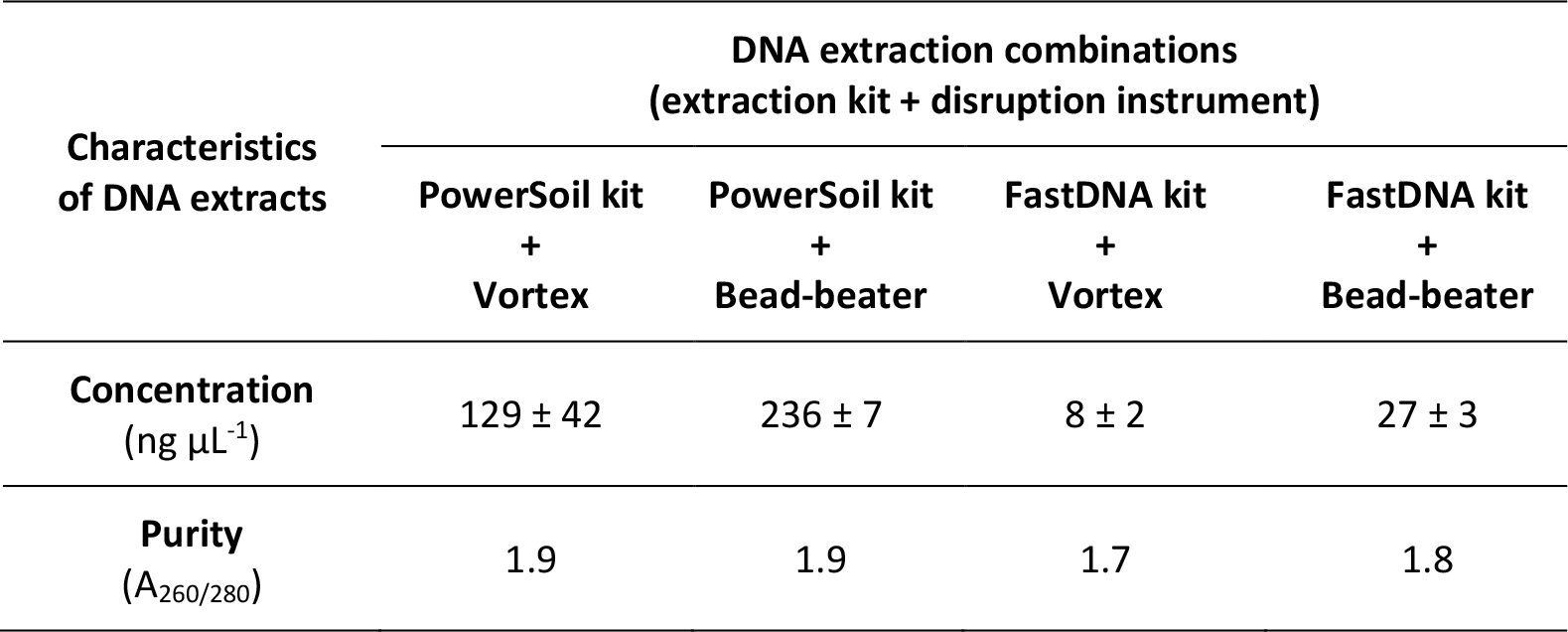
DNA concentration and purity values obtained from the four different combination of DNA extraction kits and disruption instruments. Each combination was prepared in biological triplicates from the activated sludge sample.

| Characteristics<br>of DNA extracts | DNA extraction combinations<br>(extraction kit + disruption instrument) |  |  |  |
| --- | --- | --- | --- | --- |
|  | PowerSoil kit<br>+<br>Vortex | PowerSoil kit<br>+<br>Bead-beater | FastDNA kit<br>+<br>Vortex | FastDNA kit<br>+<br>Bead-beater |
| **Concentration**<br>(ng $\mu\text{L}^{-1}$) | 129 ± 42 | 236 ± 7 | 8 ± 2 | 27 ± 3 |
| **Purity**<br>($A_{260}/A_{280}$) | 1.9 | 1.9 | 1.7 | 1.8 |

The agarose gel electrophoresis (**Figure 1**) allowed for the assessment of DNA quantity (*i.e.*, band intensity) and fragmentation (*i.e.*, distribution of band sizes). The gel confirmed the higher DNA yields with the PowerSoil kit. This was evident from the difference in band intensities. Under the imposed conditions, both the PowerSoil and FastDNA kits yielded DNA templates with lengths above 10,000 base pairs (bp), indicating relatively good DNA integrity. However, some DNA shearing was observed from the tailing of the DNA fragment length distribution in the gel. The bead-beater disruption method resulted in slightly lower DNA fragment length compared to the vortex method, when using the PowerSoil kit (10 minutes). This trend was not observed with the FastDNA kit, likely due to the shorter disruption time recommended by the protocol (40 seconds). The FastDNA kit resulted in higher maximum DNA fragment size than the PowerSoil kit, even when processed with the bead-beater. The combination of the PowerSoil kit with the bead-beater showed higher DNA yields but at the same time higher fragmentation during extraction.

**Figure 1.**
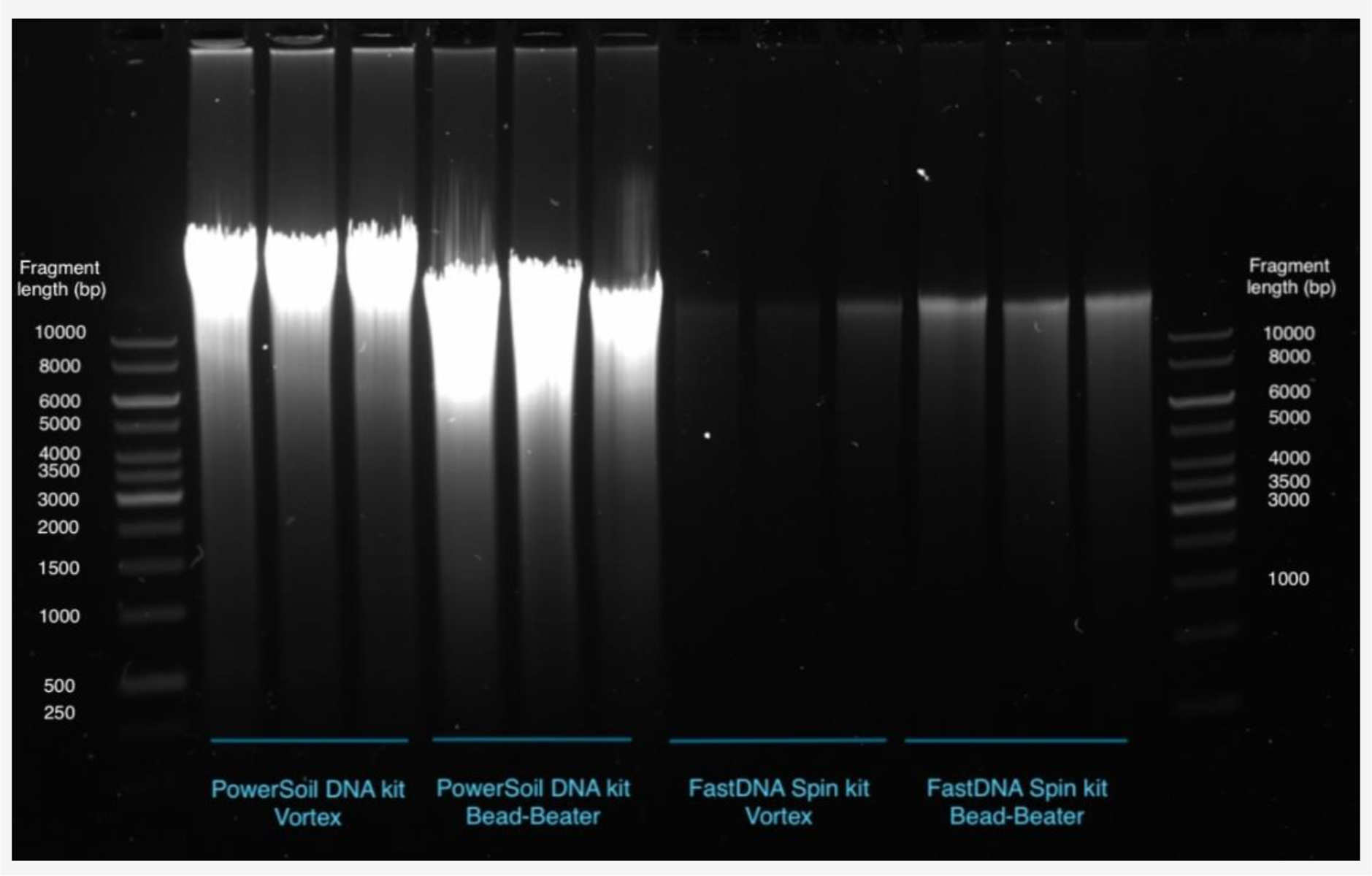
Agarose gel showing the effect of the four different combinations of DNA extraction kits (PowerSoil and FastDNA) and disruption instruments (vortex or bead-beater) on DNA yields and fragment lengths distributions. The main differences observed relate to the DNA extraction kit. Each preparation was performed in biological triplicates.

The higher DNA yield achieved with the PowerSoil kit can increase the probability of obtaining a more diverse sequencing dataset, closely representing the actual community composition due to the disruption of a larger number of cells. However, both the DNA extraction yield and integrity can influence downstream sequencing analysis. Therefore, a good quality control of DNA yield an integrity is necessary before proceeding with further molecular analysis.

### The diversity of amplicon sequencing datasets is influenced by the chosen disruption instrument and the operator factor

To assess the impact of different DNA extraction kits and disruption instruments on bacterial richness, evenness, and diversity of the amplicon sequencing datasets, we compared the alpha-diversity, Shannon index, and Simpson index (**Figure 2**).

**Figure 2.**
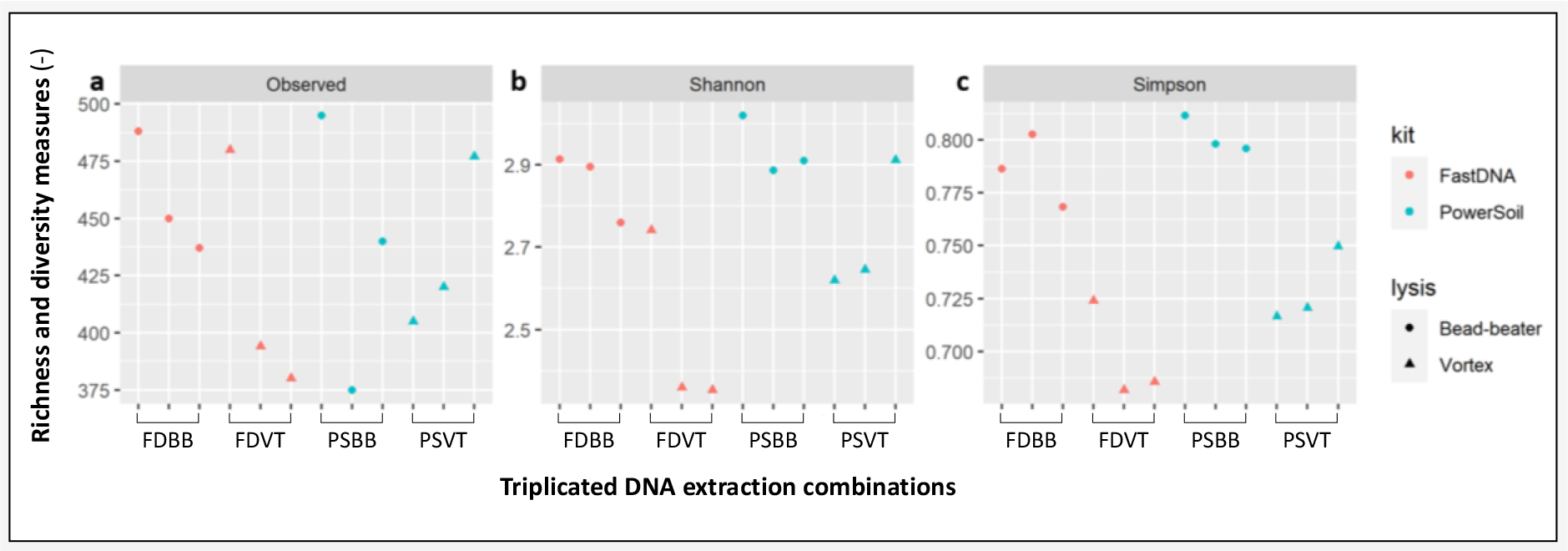
Diversity indices of the amplicon sequencing datasets obtained after the four combinations of DNA extraction kits and disruption instruments. **(a)** Alpha diversity measure from the number of observed OTUs, **(b)** the Shannon index, and **(c)** Simpson index. The number of observed OTUs, Shannon and Simpson indices were used to assess alpha diversity in all samples. Sample names are based on the DNA extraction kit and processing method. Sample code legend: FD = FastDNA Spin Kit for Soil, PS = PowerSoil, v = vortex and bb = bead-beating. Numbers represent the first, second, and third biological triplicates.

Samples extracted with vortex (*triangles* in **Figure 2**) presented lower Shannon and Simpson indices compared to those treated with the bead-beater (*circles*). Richness and evenness in the distribution of operational taxonomic units (OTUs) were lower with the vortex than with the bead-beater. The use of different DNA extraction kits did not show any significant clustering effect on alpha-diversity, Shannon index, or Simpsonindex. **Figure 2** demonstrates the impact of the operator in DNA extraction, as observed through variations between biological replicates processed by the three analysts of each group.

Bead-beating resulted in higher richness and diversity of OTUs, suggesting that the disruption mode had a greater impact than the extraction kit on the recovery of DNA from various microbial populations. The operator factor also plays a role in method harmonisation, as variations in DNA quality and yield are affected by both the analyst and the procedures followed.

### Dominant operational taxonomic units in bacterial community compositions displayed by the amplicon sequencing datasets

The observed bacterial community compositions determined based on the 16S rRNA gene amplicon sequencing datasets expressed at higher (phylum) and more specific (genus) taxonomic levels are displayed in **Figure 3** in function of the different DNA extraction treatments.

**Figure 3.**
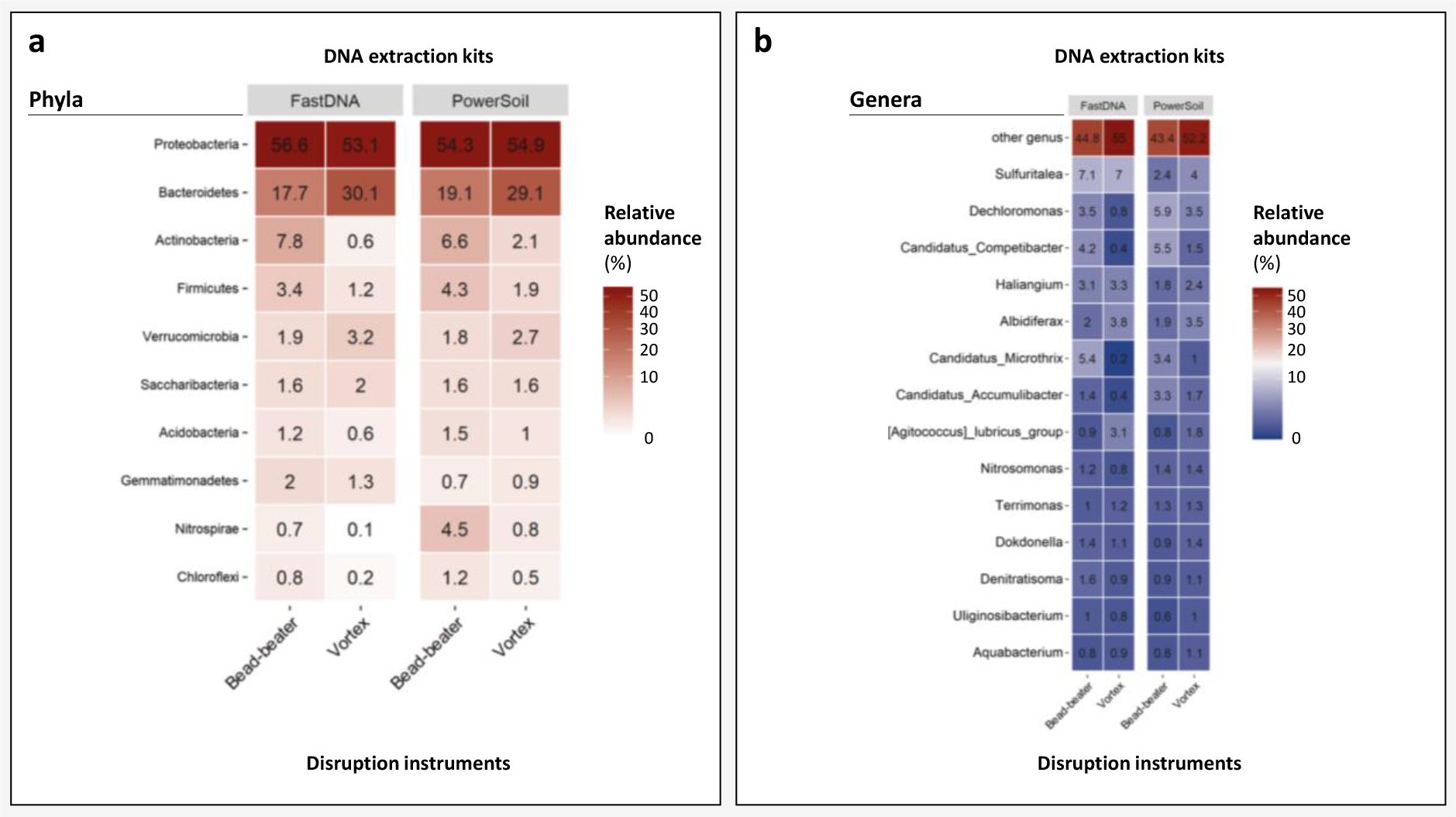
Bacterial community profiles displaying relative abundances of predominant microbial populations identified in the 16S rRNA gene amplicon sequencing datasets at phylum (**a**) and genus (**b**) levels. Heatmaps highlight the median of the relative abundance values measured from the biological replicates (see figure 6a for individual profiles). Sample code legend: sample names are based on the DNA extraction kit, processing methods and number of triplicates. Taxonomic systematics have recently been adapted (Meier-Kolthoff et al., 2022), such as for phyla where the suffix “-ota” is now implemented. The namings used in this figure referred to the previous systematics of the database used for mapping, but indications of the new names are given in the main text of the article.

As the taxonomic level becomes more specific, an increased number of unclassified OTUs were observed. The percentage of sequences categorised as “others” increased from 16-27% at the order level to 44-55% at the genus level. The heatmaps of median relative abundances over triplicates (**Figure 3**) and the bar-plots of amplicon sequencing profiles of each analyte (see **Figure 6**) produced at different phylogenetic levels (phylum and genus) revealed similar bacterial composition patterns, particularly when comparing DNA extraction kits. However, significant differences were observed among the disruption instruments.

The choice of disruption mode affected the relative abundances of specific bacterial phyla, such as *Bacteroidota*, *Actinobacteriota*, *Nitrospirota*, and *Pseudomonadota* (**Figure 3a**). It is worth noting that the taxonomic namings have recently been adapted (Meier-Kolthoff et al., 2022). Formerly, these phyla were known as *Bacteroidetes*, *Actinobacteria*, *Nitrospirae*, and *Proteobacteria*, respectively, as reflected in this figure based on the database used. For instance, *Bacteroidota* showed approximately 10% higher abundance after low-energy vortexing compared to high-energy bead-beating (30% *vs.* 18% with FastDNA and 29% *vs.* 19% with PowerSoil, respectively). This over-representation in the amplicon sequencing dataset may be attributed to potential lower cellular resistance of microbial populations belonging to this phylum to physical disruption. Underestimation of the phylum *Nistrospira* (<1%) was observed when samples were extracted with FastDNA or using the vortex, while combining PowerSoil with bead-beating resulted in a relative abundance of 4.5%. This difference between methods was much higher than the minor variations in relative abundances observed within triplicates (see **Figure 6**).

At the order level (data not shown), a similar trend was observed for the effects of vortexing and bead-beating on the order *Sphingobacteriales* (27% *vs.* 16% with FastDNA and 27% vs. 17% with PowerSoil). Combining PowerSoil with bead-beating led to higher relative abundances of *Acidimicrobiales* (2.8%), *Rhodocyclales* (1.5%), *Xanthomonadales* (2.3%), and *Clostridiales* (1.3%) compared to extractions performed by vortexing. Furthermore, the difference in relative abundance between vortexing and bead-beating for these taxa was 2 to 3-fold larger with FastDNA than with PowerSoil, except for *Clostridiales*.

At the genus level (**Figures 3b**), the FastDNA datasets showed higher relative abundance of the thiosulfate oxidiser *Sulfuritalea* (7.0% with vortex and 7.1% with bead-beater) compared to the he PowerSoil datasets (2.4% with vortex and 4% with bead-beater).

More detailed information regarding the effect of DNA extraction kits and disruption instruments on isolating core taxa can be found in the principal component analysis (PCA), which was used to visualise differences in major taxa identified from the sample groups (**Figure 4**). The bacterial community compositions of the biological triplicates processed with the same extraction methods clustered together at both phylum and genus levels. This highlights that the method combination used for DNA extraction accounted for most of the analytical variability observed in the amplicon sequencing results. However, the majority of OTUs clustered in the centre of the PCA ordinations, suggesting that, in most cases, the DNA extraction method induced slight changes in the relative abundances of microbial populations rather than their presence or absence in the datasets.

**Figure 4.**
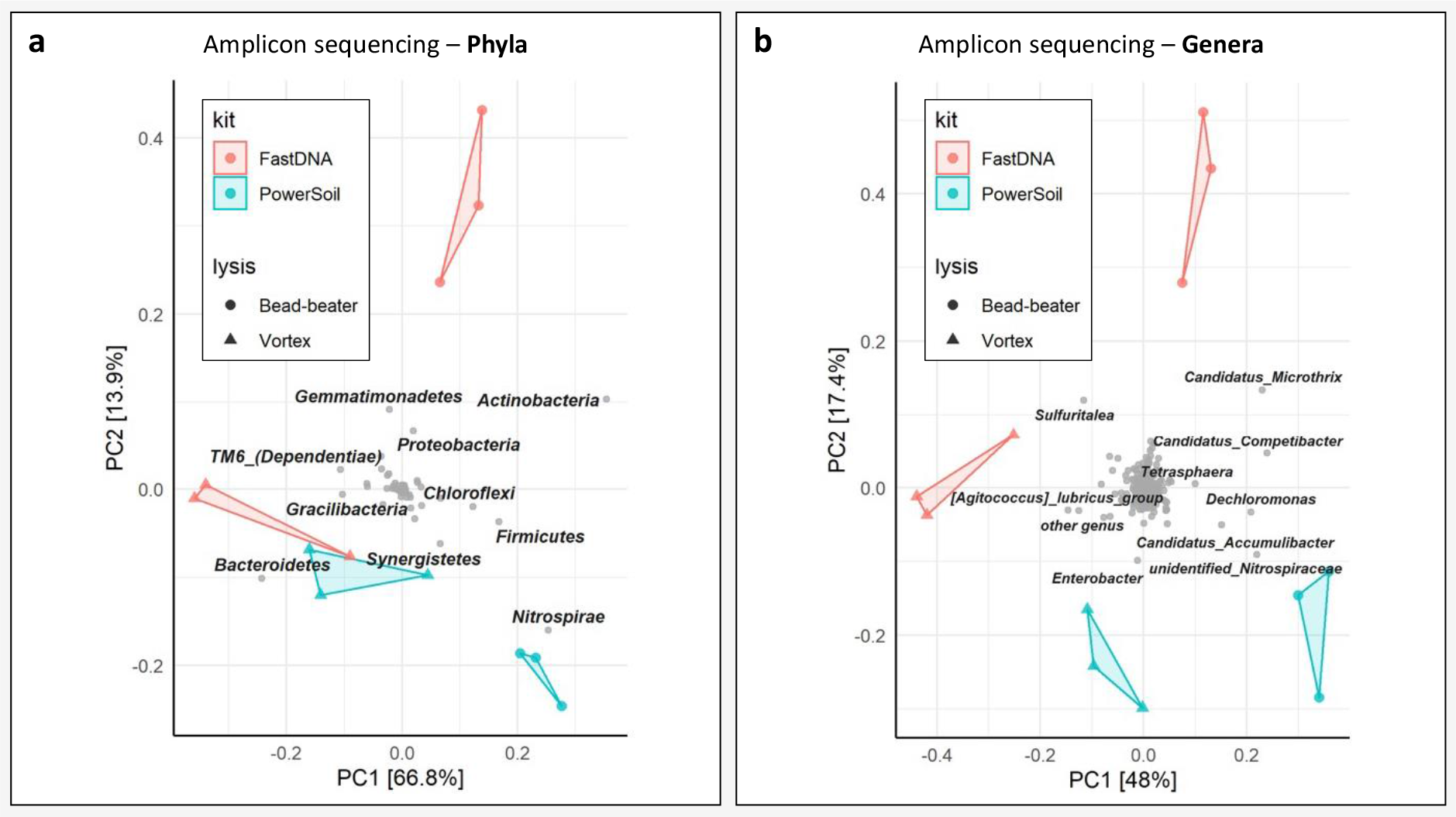
Principal component analysis (PCA) of the 16S rRNA gene amplicon sequencing datasets at phylum (**a**) and genus (**b**) levels. Prior to PCA analysis, OTU’s abundance data was obtained using the Hellinger transformation. PCA displayed the principal components and numerical differences of major core taxa between sample groups, DNA extraction (kit) and physical disruption instrument (lysis). Taxonomic identities used in this figure referred to the previous systematics of the database used for mapping, but indications of the new names of phyla are given in the main text of the article.

Nevertheless, there were some exceptions along the first component axis PC1, which explained 67 and 48% of the variability in the PCA plots at phylum and genus level, respectively. The relative abundance of the phylum *Nitrospirota* showed a significant correlation with the combined used of the PowerSoil kit and bead-beater (**Figure 4a**). Other phyla like *Bacteroidota* and *Synergistota* (formerly known as *Synergistetes*) were more prominent in the community datasets resulting from DNA extractions performed at low energy with the vortex. On the other hand, phyla like *Actinobacteriota*, *Bacillota* (formerly *Firmicutes*), and *Nitrospirota* were more represented when higher energy bead-beating was used. At the genus level, the relative abundance of “*Candidatus* Accumulibacter” and an unidentified population from the *Nitrospiraceae* family was higher when using PowerSoil (**Figure 4b**).

A consistent trend in taxonomic profiles was observed among samples processed with the same DNA extraction method. However, the disruption instruments and power, rather than the extraction kits, result in over-or under-representing specific core taxa. This effect is dependent on the disruption energy input provided during the DNA extraction process. Such considerations are important in analytical designs when investigating specific microorganisms of interest in mixed cultures, whether activated sludge or other microbial communities in environmental and health settings.

### Comparison of microbial profiles obtained by amplicon sequencing and metagenomics

In addition to DNA extraction methods, the choice of molecular analysis technique can influence the microbiome profiles obtained. This was demonstrated in this study by comparing results from 16S rRNA gene amplicon sequencing and shotgun metagenomics.

First, the application of non-metric multidimensional scaling (NMDS) analysis to the metagenomic datasets revealed distinct separation in the microbiome profiles associated with the different combinations of DNA extraction methods (**Figure 5**). This differentiation was evident when examining the data at both phylum and genus levels. The stress values (<0.1) displayed a good fit for the dimensionality reduction of the NMDS. These results align with the observations made using amplicon sequencing, further confirming that various DNA extraction method combinations introduce variability in the analysis of microbial communities.

**Figure 5.**
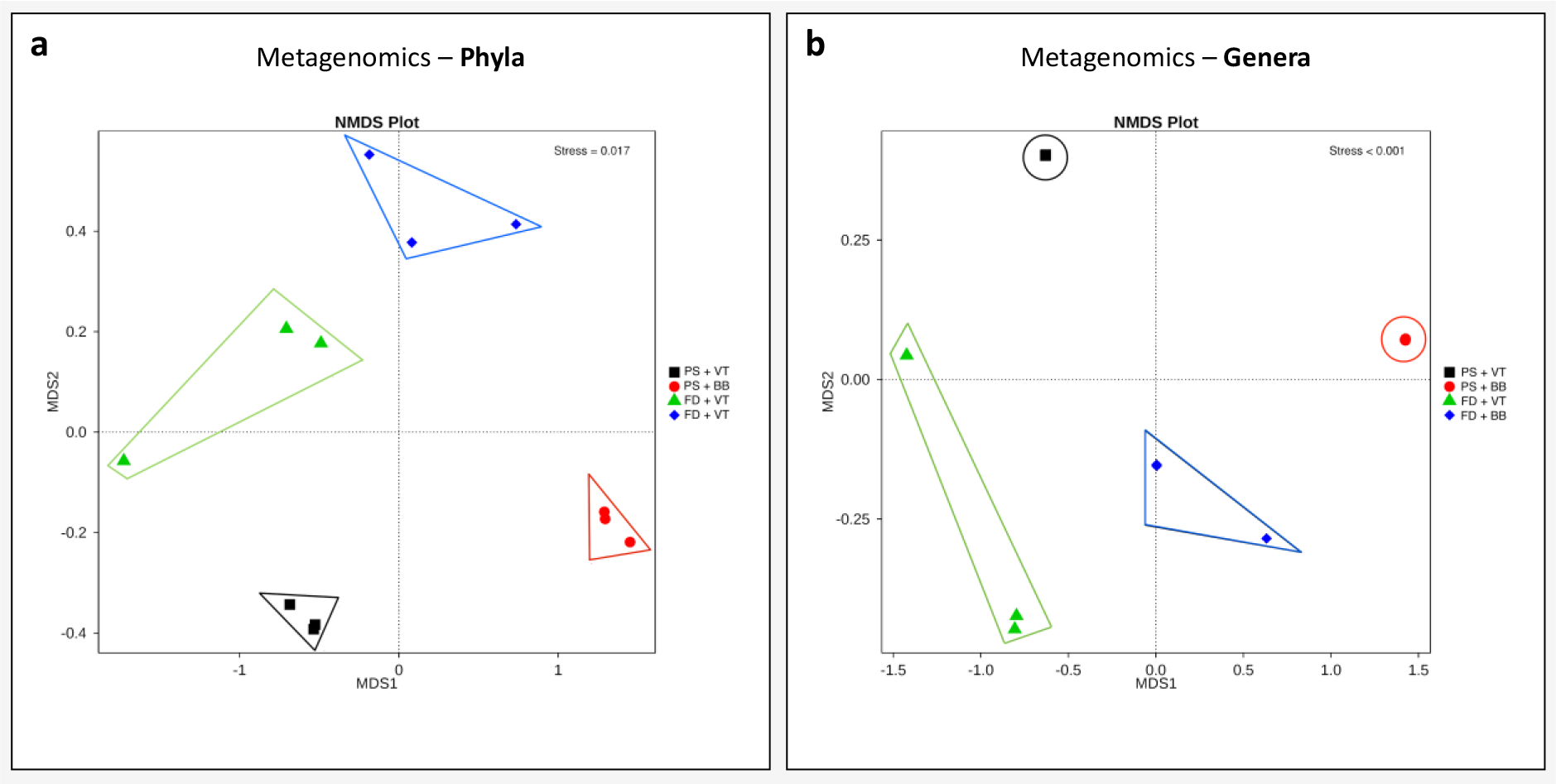
NMDS plots of the microbiome profiles computed at phylum (**a**) and genus (**b**) levels from the metagenomics datasets. Sample code legend: FD=FastDNA Spin Kit for Soil, PS=PowerSoil, VT=vortex, and BB=bead-beating.

Taxonomic classification of metagenomic reads was performed using Kraken2. The summary table presenting the number of classified and unclassified raw reads can be found in **Table S1** of the supplementary information. As is common in metagenomics, only a small fraction of reads could be effectively classified from the activated sludge datasets. The proportion of classified reads, using the standard NCBI database, ranged from 15.6% (FastDNA and vortex) to 24.3% (PowerSoil and bead-beating). When the classified reads were normalised by the total number of aligned k-mers in each sample and represented as relative abundances (**Figure 6**), some differences were observed between the classifications obtained by 16S rRNA gene amplicon sequencing (**Figure 6a**) and shotgun metagenomics (**Figure 6b**).

**Figure 6.**
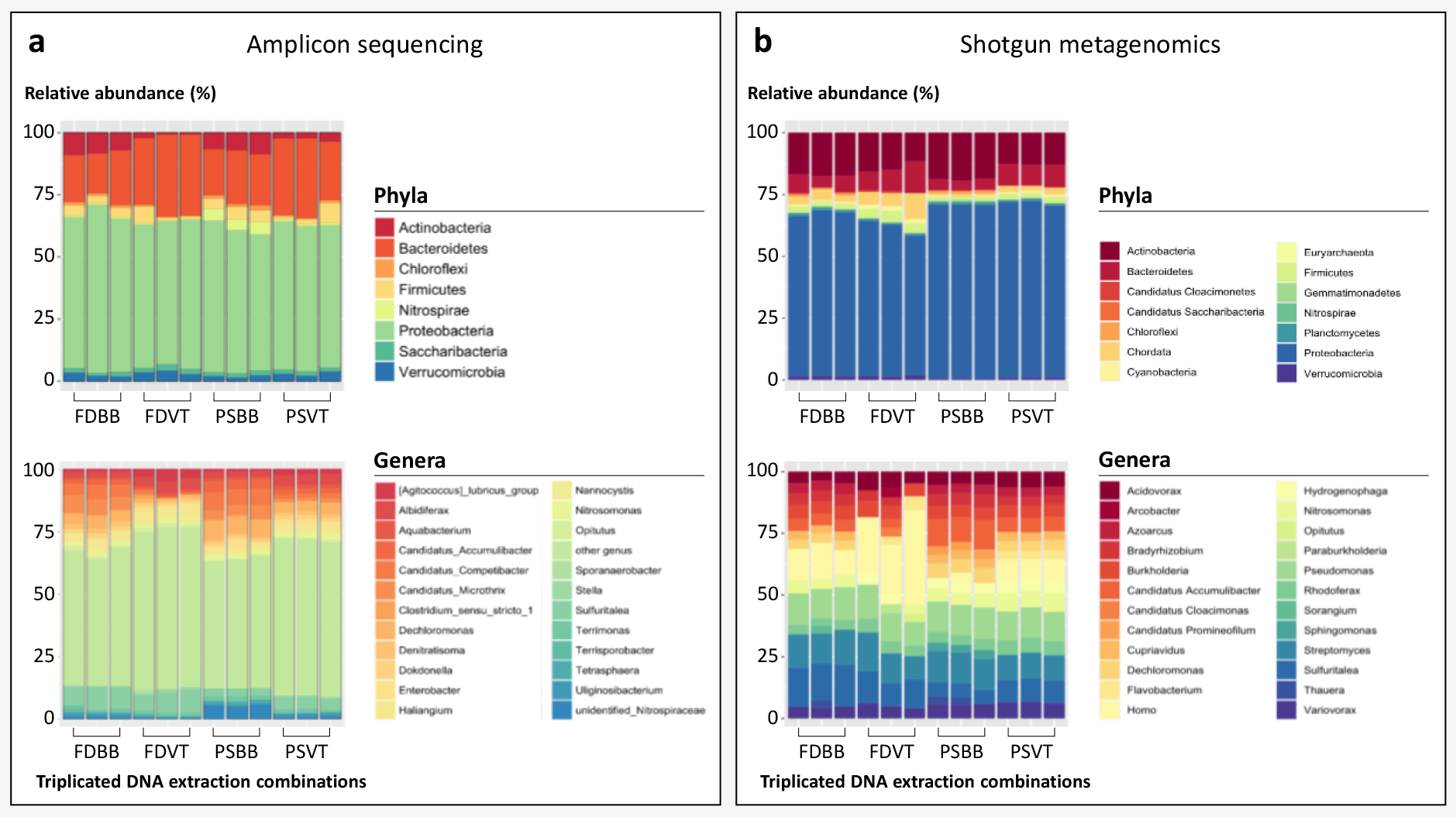
Distribution of bacterial phyla and genera in the 16S rRNA gene amplicon sequencing (**a**) and shotgun metagenomics (**b**) datasets of the activated sludge microbiome. Sample names are based on the DNA extraction kit, processing method and number of biological triplicates: FD = FastDNA Spin Kit for Soil, PS = PowerSoil, V = vortex, and BB = bead-beating. Each extraction was processed in triplicates by the four groups of operators. To compare samples, the datasets of the barplots were normalised by median sequencing depth. Only taxa that represent ≥ 1% and ≥ 1.5% of relative abundance are displayed in the phylum and genus bar plots, respectively. Taxonomic identities used in this figure referred to the previous systematics of the database used for mapping, but indications of the new names of phyla are given in the main text of the article.

A discrepancy was observed between the two sequencing approaches regarding the number of classified phyla: 16S rRNA gene amplicon sequencing identified 8 phyla, while metagenomics detected 14 phyla. The major phyla including *Pseudomonadota* (formerly *Proteobacteria*)*, Actinobacteriota, Bacteroidota, Bacillota* (formerly *Firmicutes*), and *Chloroflexota* (formerly *Chloroflexi*), were represented in similar relative abundances with both techniques. However, certain bacterial phyla like *Cyanobacteriota* (formerly *Cyanobacteria*) and *Gemmatimonadota* (formerly *Gemmatimonadetes*) could not be detected using 16S rRNA gene amplicon sequencing. The archaeal phylum *Euryarchaeota* was more easily detected by metagenomics, despite the fact that the set of PCR primers (341f and 806r) used for amplicon sequencing theoretically cover both bacteria (89% coverage of reference 16S rRNA gene sequences in ARB-SILVA according to primer testing) and archaea (83%; with *Euryarchaeota* covered at 88%) (Weissbrodt et. al, 2020).

The comparison between amplicon sequencing and metagenomics datasets became more complex when examining genera. Unclassified amplicon sequencing OTUs were categorised under “others”, while unclassified metagenomics reads were excluded from the analysis and representation. Nevertheless, the percentages of classified reads at the genus level were comparable between amplicon sequencing and shotgun metagenomics datasets (around 17-24%). At the phylum level, the community patterns were more consistent across groups and replicates when PowerSoil was used for DNA extraction. At the genus level, the number of reads classified as “others” increased in analytes obtained through vortexing, regardless of the DNA extraction kit used. Differences in the most abundant genera displayed by the two sequencing approaches were observed. Shotgun metagenomics highlighted “*Ca.* Accumulibacter”*, Pseudomonas, Streptomyces*, and *Sulfuritalea*, while amplicon sequencing displayed “*Ca.* Accumulibacter*”, “Ca.* Competibacter*”, Dechloromonas*, and an unidentified relative from the *Nitrospiraceae* family as the dominant genera.

To characterise the microbial community at lower taxonomic levels, specialised and well-annotated databases containing complete genomes should be used to increase the information from specific microbiomes such as activated sludge samples. To get the overall idea of the microbial dynamics in experimental time series or at different geographical sampling points at high taxonomic level (phylum, order, and family), amplicon sequencing provides a rapid overview. Metagenomics will have to be applied for more thorough insights on the diversity of microbial populations and functional genes, as well as on interactions between ARGs and DNA contigs affiliating with potential microbial hosts.

Amplicon sequencing provides a rapid overview for obtaining a broad understanding of microbial dynamics up to the genus level. Shotgun metagenomics should be employed to characterise microbial communities at lower taxonomic levels and to gain more comprehensive insights into the diversity of microbial populations and functional genes from microbial communities like activated sludge. This can be achieved by utilising specialised and well-annotated databases of complete genomes or near-complete metagenome-assembled genomes (MAGs). In addition to microorganisms, metagenomics help unravel the pool of ARGs carried by the community and its microbial populations.

### qPCR to assess DNA extraction artifacts

qPCR was conducted on the two ARGs *sul1* and *bla*_CTX-M_, which represent two classes of antibiotics, (sulfonamides and betalactams) that are commonly consumed in The Netherlands (Pallarés-Vega et al., 2020). The variations in the measured amount of specific ARGs per mg of fresh activated sludge were primarily attributed to the disruption instrument rather than the DNA extraction kit (**Figure 7**). This matched with instrumental effects on the 16S rRNA gene amplicon sequencing patterns.

**Figure 7.**
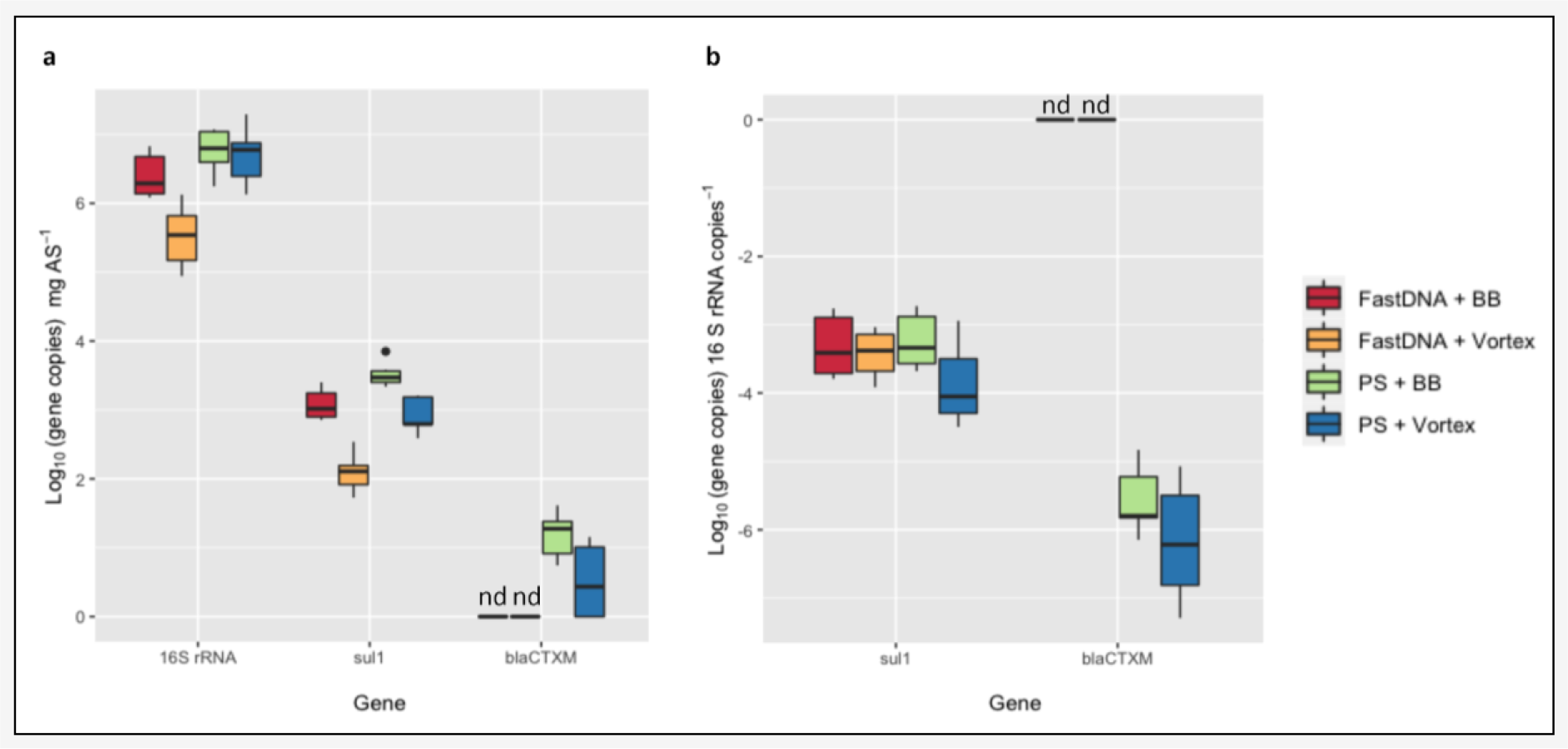
Comparison of the qPCR analysis of 16S rRNA gene and the selected ARGs from the DNA fractions of activated sludge isolated from the combination of the different methods. Values are displayed as log_10_ (gene copies) mg^−1^ of fresh activated sludge (**a**) and normalised by the *16S rRNA* gene (**b**). The “nd” labels indicate gene not detected.

However, in cases where the abundance of a specific ARG like *bla*_CTX-M_ was low, the DNA extraction kit played a significant role. In this instance, *bla*_CTX-M_ was not detected in the DNA samples extracted with FastDNA, regardless of the disruption instrument used.

Furthermore, the vortexing of FastDNA kits resulted in a lower number of gene copies compared to the other combinations. This went along with a significant difference of 0.87 ± 0.22 log_10_ (gene copies) for the 16S rRNA gene and of 0.98 ± 0.14 log_10_ (gene copies) for the *sul1* ARG, per mg of activated sludge (*p* < 0.005) (**Figure 7a**). Since *bla*_CTX-M_ was not detected in DNA pools extracted with FastDNA, a similar comparison could not be performed for this gene. When ARGs were normalised by 16S rRNA gene copies, no differences were observed, indicating that the DNA extraction kit and disruption instrument equally affected the detection and quantification of both the reference 16S rRNA gene and the ARGs (**Figure 7b**).

Before a detailed analysis of the resistome from a microbial community like activated sludge, the analysis of selected genes by qPCR provides a rapid first overview of the influence of different DNA extraction workflows on molecular results at both bacterial and ARG level.

### Effect of DNA extraction kits and disrupting instruments on resistome analysis

The analysis of the 12 metagenomes revealed the presence of about 145 genes associated with AMR in microorganisms present in the activated sludge (**Figure 8a**). These ARGs, as classified in the CARD database, belonged to 49 different gene families. Among these, 95 genes confer resistance to single antibiotic groups, 5 genes to another antimicrobial group (*e.g.*, Triclosan), and the remaining 45 genes were associated with resistance to 2 to 10 different antibiotic groups.

**Figure 8.**
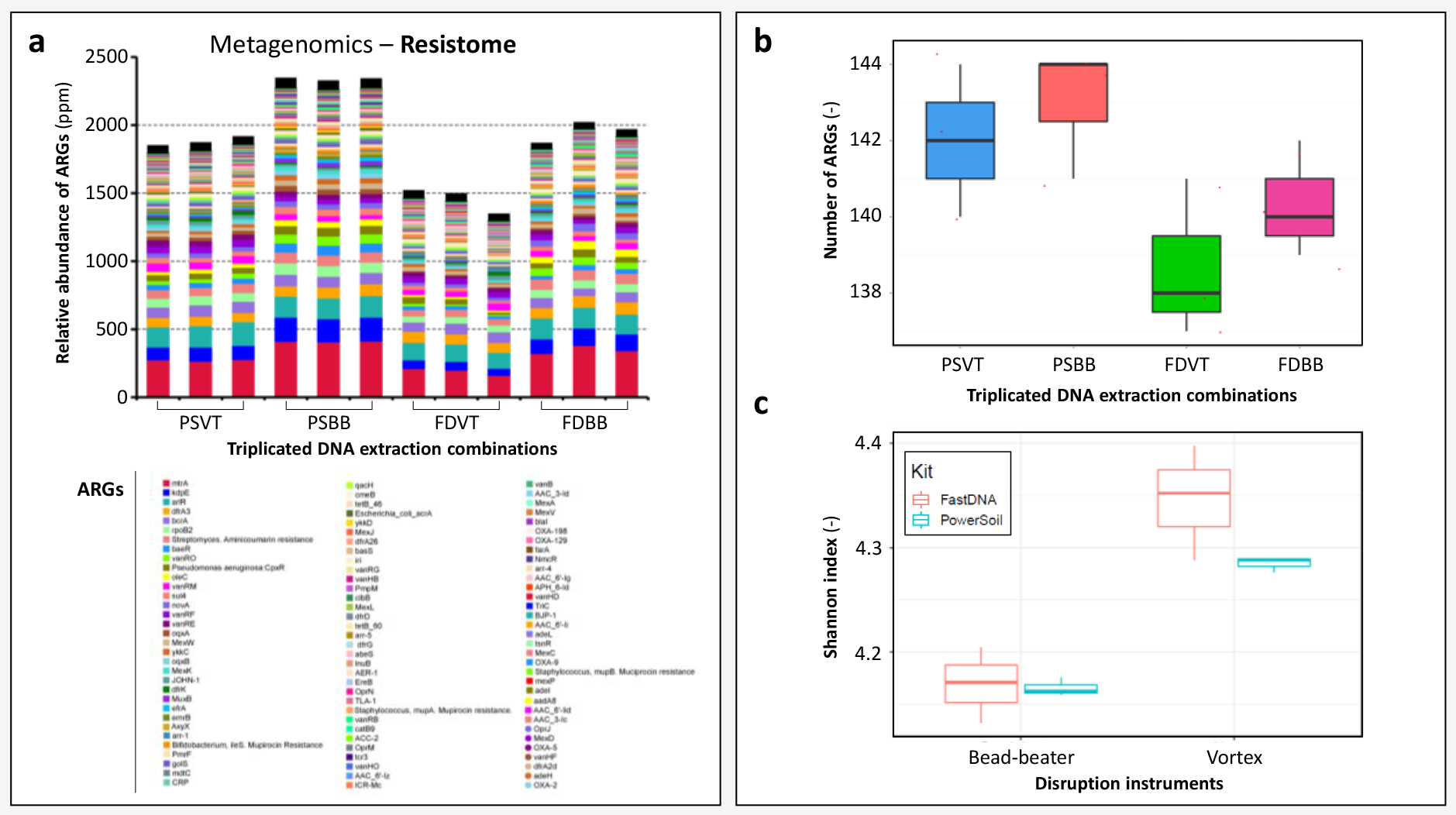
Resistome profiles measured by metagenomics from the 12 DNA extracts obtained from the triplicated combinations of DNA extractions performed on the activated sludge sample (**a**): FD = FastDNA Spin Kit for Soil, PS = PowerSoil, V = vortex, and BB = bead-beating. Effect of the DNA extraction kits and disruption instruments (**b**) the number of antibiotic resistance genes (*i.e.*, richness of ARGs) and Shannon diversity of ARGs calculated from their absolute abundances (**c**). Each analysis was performed as biological triplicates.

When evaluating the number of identified ARGs, only negligible differences were observed the various DNA extraction kits and disruption instruments. The detected genes ranged from 138 ± 2 ARGs when DNA was extracted by vortexing the FastDNA kit to 142 ± 2 ARGs by bead-beating the PowerSoil kit (**Figure 8b**).

Despite the minimal variation in the number of observed ARGs, differences were evident when assessing the diversity of these genes based on the disruption instruments used. This was evaluated using different metrics such as the Shannon diversity index (**Figure 8c**). The overall regression analysis showed a significant fit (R^2^ = 0.85, F(3, 8) = 21.59, p < 0.05). There was not a significant effect of the DNA extraction kit used (β = −0.003, p = 0.89), but the disruption instrument had a significant effect (β = 0.18, p < 0.05). The interaction between the DNA extraction kit and disruption instrument was not significant (β = −0.06, p = 0.17). The use of the bead-beater as a disruption instrument increased the number of observed ARGs, while the FastDNA kit resulted in more variability in the diversity of genes detected by the different groups of analysts.

In addition, the cluster heatmap in **Figure 9** presents the hierarchical effects of the disruption instruments (bead-beater and vortex) and DNA extraction kits (Powersoil and FastDNA) on the detection and relative abundance of the predominant ARGs detected in the resistome of the activated sludge. The ARG abundance data are primarily discriminated by the type of disruption instrument, before the type extraction kit used. All biological triplicates conducted through the DNA extraction combination of each group clustered together. Bead-beating resulted in greater relative abundances of the ARGs. Bead-beating the PowerSoil kits during 10 minutes resulted in the largest relative abundances, while vortexing the FastDNA kits during 40 seconds led to lowest relative abundances. When following the manufacturers’ instructions (*i.e.*, vortexing of PowerSoil during 10 min and bead-beating of FastDNA during 40 seconds), the overall detection levels were similar with local differences depending on specific genes. Notably, genes coding for resistance to the vancomycin drug of last resort (*vanRM*, *vanRF*, *vanRE*) were better extracted with the PowerSoil procedure.

**Figure 9.**
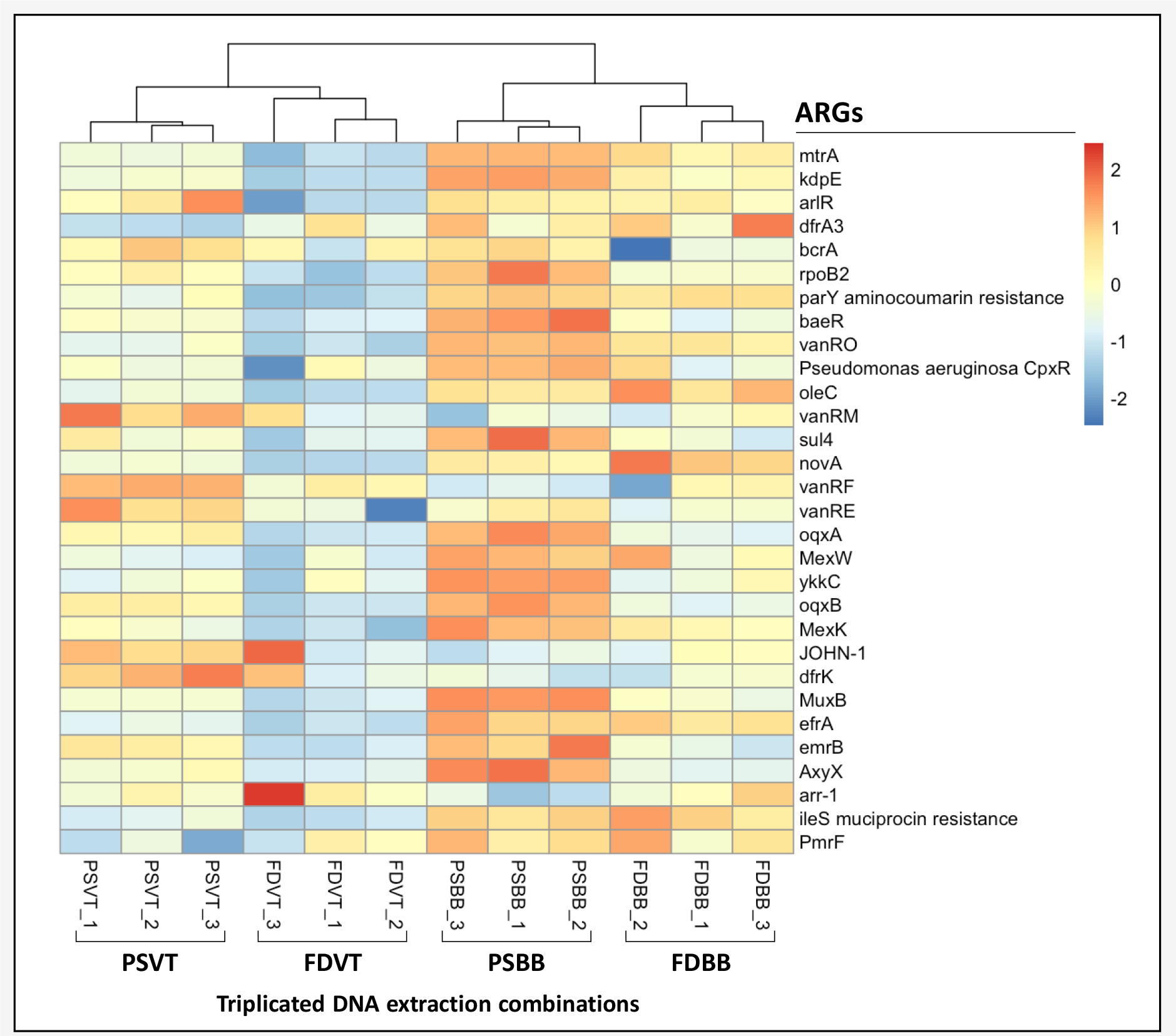
Clustered heatmap of predominant ARGs detected from the metagenomes of the 12 DNA extracts obtained from the four triplicated DNA extraction combinations: FD = FastDNA Spin Kit for Soil, PS = PowerSoil, V = vortex, and BB = bead-beating.

## Discussion

The increasing utilisation of amplicon sequencing and metagenomics in studying the microbiome and resistome of wastewater environments has emphasised the necessity for more testing of DNA extraction protocols. Harmonisation in molecular methods is crucial to ensure reliable comparisons of results across different studies conducted in various laboratories with different equipment. Our findings highlight that DNA extraction procedures have the most significant influence among the variables involved, including environmental sample type, PCR conditions (when applicable, *e.g.*, for amplicon sequencing and qPCR), sequencing machine, algorithms for assembling, binning and annotating sequencing reads and contigs, databases, and bioinformatics expertise level.

Previous studies have already demonstrated the impact of DNA extraction parameters on microbial community profiles obtained by 16S rRNA gene amplicon sequencing (Albertsen et al., 2015). The effect of the sample types has also been examined in various contexts, ranging from activated sludge (McIlroy et al., 2009; Vanysacker et al., 2010; Guo and Zhang, 2013; Albertsen et al., 2015) to human samples (Sui et al., 2020), soil (Martin-Laurent et al., 2001), and pathogenic bacteria (Bushon et al., 2010), among others. The effects of biomass sample preservation and DNA extraction methods have also been studied (Li et al., 2018).

Currently, two of the most widely used and recommended commercial kits for DNA extraction from activated sludge samples are the FastDNA Spin kit for Soil and the PowerSoil kit. Existing literature has thus far reported that extracting DNA by bead-beating FastDNA kits yields optimal results in terms of DNA yield and purity, and taxonomic profiling of activated sludge (Vanysacker et al., 2010; Guo and Zhang, 2013; Albertsen et al., 2015; Li et al., 2018). However, these results have been obtained often using the bead-beater sold by the manufacturer of kits, which may not be available in all laboratories.

Here, we provided the first systematic assessment of the influence of the combination of these two DNA extraction kits that are frequently used in practice in combination with bead-beating or vortexing, on the DNA extraction efficiency from activated sludge and the profiling of its underlying microbiome and resistome by amplicon sequencing (bacterial community profile), metagenomics (microbiome and resistome), and qPCR (quantification of selected ARGs).

### The importance of extraction kits and disruption modes on DNA yield, purity, and integrity

We obtained a higher DNA yield and purity when DNA was extracted with the PowerSoil kit compared to the FastDNA kit. The use of FastDNA resulted in a lower fragmentation of DNA than PowerSoil. A trade-off between DNA yield, purity, and integrity should be found when optimising DNA extraction protocols. Efficiencies in cell lysis in DNA extraction tubes have a significant effect on DNA recovery from microbial biomass prior to DNA elution from the adsorption columns (Martin-Laurent et al., 2001; Guo and Zhang, 2013; Li et al., 2018). Vigorous mechanical homogenisation methods efficiently lyse microbial cells to release DNA, but prolonged disruptions at high energy unfavourably shear the DNA. The duration of the cell disruption process recommended by manufacturers (10 minutes with PowerSoil, 40 seconds with FastDNA) was the main factor impacting DNA extraction, matching with earlier reports (Albertsen et al., 2015). However, authors have also observed a decline in DNA integrity after bead-beating series of 4⊆40 seconds (*i.e.*, 160 seconds), which is still much lower than the 10 minutes applied to PowerSoil. These times were applied to both the bead-beater and vortex using these kits. With FastDNA, vortexing for 40 seconds resulted in lower performance compared to bead-beating for the same duration. But the DNA yields obtained with this kit and the Mini-Beadbeater-24® were still lower than those reported with the homogeniser recommended by the manufacturer.

In general, DNA extraction kits should be used together with the instrument provided by the same manufacturer. According to protocols, FastDNA kits are recommended to be bead-beated on a FastPrep® instrument. PowerSoil kits are recommended to be used with a common bench-top vortex; but some labs also use it with a bead-beater. Not all laboratories are equipped with the specific homogenisers, and several groups use alternatives to disrupt samples. We investigated whether laboratories equipped with either high-energy homogenisers (here, Mini-Beadbeater-24®) or low-energy vortexes (here, Vortex-Genie 2®) could achieve comparable DNA extractions, while using the disruption times recommended by the manufacturers of the kits. Higher DNA yields were obtained using bead-beating, a high-energy method, compared to vortexing, a low-energy method, with both PowerSoil (1.8-fold difference in yields between bead-beating and vortexing) and FastDNA (3.5-fold difference). A short disruption time of 40 seconds can therefore be sufficient to effectively lyse the cells by bead-beating. Vortexing is a gentler method that deploys much lower energy and longer times of 10 minutes are needed to disrupt cells. Therefore, the PowerSoil kit has been recommended by its manufacturer to be used for 10 min but using a vortex. A much shorter disruption period should be implemented when using this kit with a bead-beater, to prevent a pronounced shearing of the DNA.

Combinations of DNA extraction protocols and disruption methods do not lead to equal results in biomolecular profiles, as observed in amplicon sequencing, metagenomics, and qPCR analyses. A trade-off should be found between the costs of the instruments and the quality of the data. The initial investment in the optimal combination of extraction kits and disruption instruments will offer long-term reliability in processing large sets of samples. Depending on the available disruption instrument and the type of biomass and cells to disrupt, DNA extraction protocols can further be adapted with various options. These possibilities include: a longer bead-beating time; several DNA extractions in sequence from the same tube; repetition of thermal-mechanical sequences (heat-disrupt-cool on ice); flash-freezing of biomass in liquid nitrogen prior to bead-beating; fragilisation of cells with lysozyme; or a combination of these methods.

In addition, the processing of large amounts of samples can benefit from the use of a DNA extraction robot (Weissbrodt et al., 2012; 2014). The implementation of DNA extractions at WWTP lines or even online on robotic devices aimed to collect, process, and analyse environmental samples on the collection sites necessitate the adaptation of DNA extraction workflows, since bead-beating may not be possible in every situation. When modifying protocols, it is important to always consider the optimum in DNA yield, purity, and integrity as endpoint. We strongly recommend conducting a preliminary testing, like what we performed in this study, for each kit, disruption device, and sample type that are intended to be used prior to processing the whole set of samples collected during a campaign or experiment.

Other parameters like the mass of activated sludge used in the extraction tubes and the sizes of the beads of the lysing matrix present in the tubes are frequently suggested as additional factors to consider. For instance, Li et al. (2018) have obtained a high DNA yield when extracting DNA from activated sludge samples at 2.4-3.1 g TSS L^-1^ when using FastDNA. Here, the activated sludge comprised 4.5 g TSS L^-1^, but the four groups normalised the biomass input in extraction tubes to 250 mg of centrifuged wet biomass, such as recommended by manufacturers, but mainly for soil matrices. However, activated sludge contains a much higher biomass content than a soil sample. Overall, the mass of biomass placed in extraction tubes should be considered and normalised across all extractions to ensure consistency with the nominal amount that can be extracted using the manufactured kit.

The protocol of the MiDAS Field Guide of microorganisms of wastewater treatment plants (McIlroy et al., 2015) suggests a mass of total solids equivalent to a dry weight of 2 mg (1-4 mg) to insert in the extraction tube. his mass can be achieved by weighing on a high-precision microbalance (which can be challenging because of the low mass), or by transferring a volume of biomass sample of known concentration by pipetting using wide micropipet tips (to prevent clogging of tips) to an equivalent amount of 2 mg. This can be done after homogenising the biomass with a Potter-Elvejhem tissue grinder mounted on a rotor (Weissbrodt et al., 2014), prior to centrifuging, resuspending, and washing it with volumes of a phosphate saline buffer equal to the initial volume of the mixed liquor. For an activated sludge concentration of 4.5 g TSS L^-1^, a volume of washed, homogenised mixed liquor of around 450 μL will be needed to insert 2 mg in the DNA extraction tube. A good care should be observed with respect to the maximum volume capacity of the kit. Depending on the situation, the biomass can either be diluted with phosphate-buffered saline (PBS) if the volume to pipet in the extraction tube is too low, or concentrated by spinning it down prior to resuspending it in a lower volume of PBS if the volume to pipet is too high. Better than total solids, the content of volatile solids of the biomass should be measured and considered since inorganic solids make up to 10-40% of the total solids of the activated sludge depending on process conditions, but this needs one additional day of preparation.

### Are DNA extracts representative of all microorganisms present in activated sludge?

Incomplete disruption instruments have already been shown to be an issue for the analysis of bacteria and fungi in environmental samples (Starke et al., 2019). Different DNA extraction techniques can pose a potential issue of underrepresentation in the detection of ARB. No DNA extraction method is suitable for all kinds of microorganisms since their susceptibility towards different lysis methods highly vary (De Bruin and Birnboim, 2016). The same conclusion has been formulated regarding the suitability of DNA extraction methods for different microorganisms when analysing human microbiota samples by metagenomics (Yuan et al., 2012).

Our results match with these previous studies. The efficiency of DNA extraction methods can be compared based on the presence of harder-to-lyse Gram-positive bacteria such as *Actinobacteriota* and *Bacillota* (formerly *Firmicutes*). Their thick cell wall or the ability of some of them to form spores, provide high resistance against chemical and physical disruption (Guo and Zhang, 2013; Albertsen et al., 2015). Over- and underrepresentation of specific taxa in the resulting DNA samples should be acknowledged when aiming at extracting DNA from targeted microorganisms using different disruption instruments. DNA extraction protocols are often intentionally adapted by laboratories for specific needs. The aforementioned adaptations can improve the extraction of DNA from these hard-to-lyse cells, which will also impact the resulting community profile. Mixtures of a representative set of known microbial populations (*i.e.*, mocked communities made of isolates or enrichments) can be used as standards for comparing DNA extraction efficiencies across studies.

### Increasing analytical resolution and sensitivity by combining molecular methods

#### Benefits and challenges of amplicon sequencing for rapid community overviews

The 16S rRNA gene has long been utilised as a universal marker for bacterial community analysis. 16S rRNA gene amplicon sequencing provides valuable insights into the composition of bacterial populations, allowing for the investigation of temporal and geographical dynamics in both technical and natural environments (Cerruti et al., 2021).

However, the accuracy of microbial community profiling can be influenced by biases introduced during DNA extraction and PCR protocols, leading to misrepresentations of specific microbial populations. Another crucial factor contributing to analytical differences in community compositions is the choice of the hypervariable region of the 16S rRNA gene (Gohl et al., 2016; Rausch et al., 2019; Weissbrodt et al., 2020). Additionally, the presence of multiple copies of the 16S rRNA gene in bacterial genomes further complicates the interpretation of community profiles. While correction factors have been developed for known microorganisms with reference genomes, this is not feasible for unknown microorganisms. Nevertheless, OTUs are commonly defined at a 97% similarity threshold, which also helps mitigate the impact of 16S rRNA gene copies within an operon. The influence of multi-copies can become more significant when amplicon sequence variants (ASVs) are used for profiling instead of OTUs. Notably, multi-copies mainly affect the normalisation of qPCR data.

To overcome this, the utilisation of the single-copy *rpoB* gene as emerged as a viable alternative for qPCR (Case et al. 2007; Calderón-Franco et al. 2022). However, the use of *rpoB* as an amplicon sequencing marker is marginal due to the lack of a comprehensive database of reference sequences of this gene across the entire microbial tree of life, as well as the non-universality of *rpoB* sequences *per se*. Exploring the feasibility of extracting *rpoB* genes from MAGs may hold promise for future biomarker studies.

#### Expanding microbial diversity assessment and ARG profiling using metagenomics

Shotgun metagenomics offers several advantages over targeted approaches, providing a higher resolution for assessing the diversity of prokaryotes, eukaryotes, and viruses. Together with the binning and annotation of MAGs from single lineages within a microbial community (Albertsen et al., 2013; Park et al., 2017), it also enables strain-level classification of microorganisms, unveiling a wealth of unexplored microbial life that would otherwise remain unclassifiable. Moreover, it empowers researchers to investigate the functional relationships between hosts and bacteria by directly examining the functional genetic content of samples.

However, database improvement is crucial as a significant portion of reads (>75%) remains unclassified. This could be attributed to unknown microbial populations and limitations in database annotation, such as sequence duplications or incomplete annotations. Collaborative efforts, exemplified by the MiDAS Field Guide to the Microbes of Activated Sludge and Anaerobic Digesters (Dueholm et al. 2022), are resulting in the development of ecosystem-specific unified databases containing long-read 16S rRNA gene sequences and high-quality MAGs from wastewater microorganisms worldwide (Singleton et al., 2021). The annotation and integration of MAGs into specialised databases are essential for comprehensive classification and detailed study of microorganisms, including their functional potential. ARGs annotated from high-quality MAGs have been recently integrated in the MiDAS database (https://www.midasfieldguide.org/guide/antibiotic-resistance). Integration of these functional genetic signatures with other multi-omics methods can shed light on the expression and regulation of metabolisms, such as antibiotic resistance phenotypes.

Metagenomics also provides valuable insights into the diversity of functional genes present in microbial communities, including ARGs in the resistome of activated sludge. Through metagenomics, a set of 145 ARGs was identified in the activated sludge at the sequencing depth used. Furthermore, metagenomics allows for the association of ARGs with specific microbial populations, including potential pathogens, by analysing both functional genes and taxonomies from genetic contigs. This analysis can also reveal the co-localisation of ARGs on mobile genetic elements (MGEs) like plasmids, which facilitate the transfer of ARGs bewteen microbes (Calderón-Franco et al., 2022). Innovative sequencing techniques like Hi-C metagenomics, which leverages proximity ligation of MGEs with genomic DNA prior extraction and sequencing, can provide insights into which specific populations within the microbial community harbour or have acquired specific MGEs and ARGs in their accessory genome (Calderón-Franco et al., 2023). This approach can considerably enhance our understanding of lateral gene transfer and acquisition processes in mixed microbial cultures.

Although the number of ARGs obtained showed minor differences among the different DNA extraction strategies, the disruption instrument used can influence the observed amount of ARGs. This discrepancy can be attributed to the high disruption energy achieved with bead-beating compared to vortexing, resulting in a higher yield of DNA that can be purified and sequenced. Consideration of the disruption instrument is therefore crucial when assessing ARG diversity, as reflected by the Shannon index.

Overall, shotgun metagenomics provides a powerful tool for unravelling microbial diversity, ARG and functional gene content, and their interactions with microorganisms, while database improvements and innovative techniques will enhance our understanding of microbial ecosystems and their role in various processes like AMR spreading.

#### Metagenomics and qPCR are complementary for high resolution and sensitivity in AMR studies

When it comes to characterising ARGs in AMR studies, the question often arises as to whether metagenomics or qPCR should be used. The answer lies in utilising both approaches. Metagenomics provides valuable insights at high resolution into the diversity of genes and the populations potentially carrying these genes. qPCR offers higher sensitivity in detecting specific target genes of interest.

The combination of metagenomics and qPCR has demonstrated significant benefits in AMR studies, particularly in wastewater environments. This approach enables comprehensive profiling of microbiomes and resistomes, shedding light on their compositions and dynamics at high resolution and sensitivity (Calderón-Franco et al., 2022). Recent advancements in highly-multiplexed qPCR, high-throughput qPCR, and smart chips have made it possible to simultaneously measure hundreds of genes. This enhanced sensitivity allows for precise tracking of genes and their prevalence in water samples, facilitating the development of early-warning systems for environmental and health issues through gene surveillance campaigns (Lai et al., 2021).

#### Ensuring accurate DNA extractions in microbiome and resistome studies

It is crucial to give careful attention to DNA extractions in microbiome and resistome studies. Neglecting this step can introduce biases that affect downstream analytical methods, including qPCR, multiplex PCR, amplicon sequencing, metagenomics, and Hi-C sequencing among others. These biases can hinder accurate characterisation of genetic compositions and processes in microbial communities of wastewater environments. This concern extends to any molecular and multi-omic analysis involving the extraction of informational macromolecules such as DNA, RNA, proteins, lipids, and polysaccharides, which are inherently stable structural components of microbial cells and not naturally easily extractable.

To advance beyond local, purpose-driven implementation of methods, the development of harmonised protocols across groups becomes crucial. These protocols enable inter-comparability, reproducibility, and the design of larger collaborative studies to address a wider range of analyses related to AMR determinants and associated environmental and health problems (Miłobedzka et al., 2022). DNA extraction is an integral part of this process and requires special attention to ensure accurate and reliable results. The importance of educating the next generations of scientists on these endpoints, as demonstrated in this work, cannot be overstated.

## Conclusions

We conducted this study to learn and investigate the influence of DNA extraction methods on the analysis of activated sludge using various techniques such as 16S rRNA gene amplicon sequencing, shotgun metagenomics, and qPCR. Our findings revealed that the choice of DNA extraction method influence the profiles of bacterial communities, microbiomes, and resistomes obtained through these molecular analyses. It emphasised the need for harmonising protocols to facilitate the comparison of molecular datasets across different research groups. This study contributes to international efforts in harmonising protocols for analysing microbial communities and AMR determinants in wastewater. It also highlights the importance of understanding the limitations and challenges associated with molecular methods in wastewater microbiology and engineering. The incorporation of this approach into scientific projects and practical courses in environmental molecular biology, along with training the next generation of experts, is an important target for a better management of AMR spreading in water systems.

## Author contributions

DCF, DK, BA, and DGW organised the workshop and designed the study. DCF, DK, BA, JAS, JR, RPV, MA, SA, SG, IK, MS, and SP performed the wet-lab preparations in 4 groups of 3 analysts. DCF, DK, RD, MALM, and DGW analysed the data. DCF, DK, and DGW prepared the manuscript, with inputs and edits by the contributors.

## Acknowledgements

This study was funded by the EU H2020 Twinning project REPARES – Research Platform on Antibiotic Resistance Spread Through Wastewater Treatment Plants (https://repares.vscht.cz/). It resulted from a REPARES workshop organised at the TU Delft in The Netherlands with various project partners, aimed at educating young researchers on molecular methods for wastewater microbiology and engineering, as well as the influence of DNA extraction workflows on analytical outputs.

This manuscript is deposited as a pre-print in *bioRxiv*.

## Supplementary information

**Figure S1.**
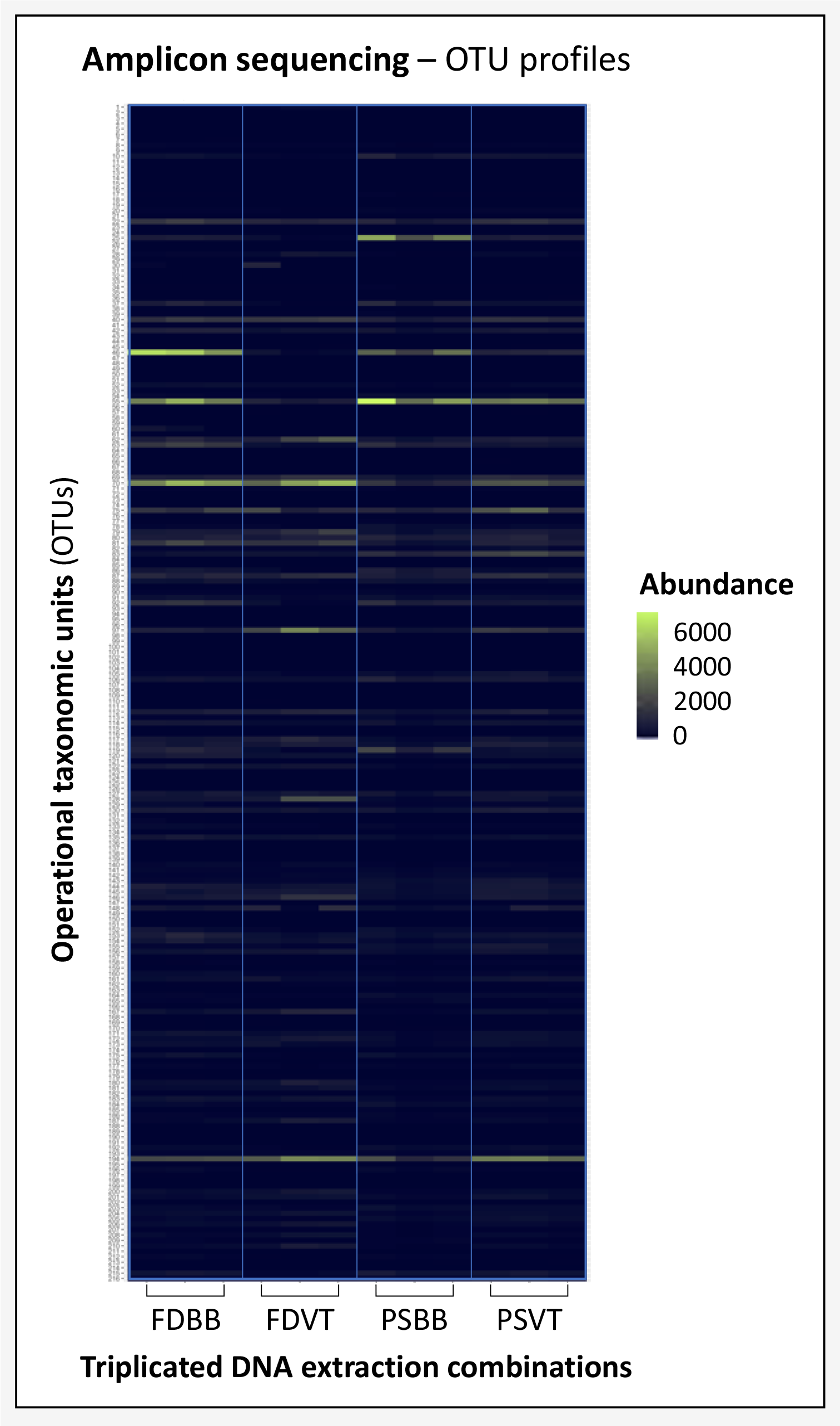
Heatmap of abundances of operational taxonomic units (OTUs) detected from the 16S rRNA gene amplicon sequencing datasets obtained according to the different combinations of DNA extraction kits and disruption instruments: FD = FastDNA Spin Kit for Soil, PS = PowerSoil, V = vortex, and BB = bead-beating.

**Table S1.** Summary table of the metagenomics classification obtained from Pavian.

| Name | Number of raw reads | Classified reads | Chordate reads | Artificial reads | Unclassified reads | Microbial reads | Bacterial reads | Viral reads | Fungal reads | Protozoan reads |
| --- | --- | --- | --- | --- | --- | --- | --- | --- | --- | --- |
| FDBB_1 | 22,347,514 | 18.8% | 0.528% | 0% | 81.2% | 18.2% | 18% | 0.024% | 0% | 0.0000224% |
| FDBB_2 | 26,461,180 | 21.1% | 0.812% | 0% | 78.9% | 20.2% | 20% | 0.0321% | 0% | 0.0000189% |
| FDBB_3 | 21,756,666 | 19.6% | 0.436% | 0% | 80.4% | 19.2% | 18.9% | 0.0228% | 0% | 0.0000322% |
| FDVT_1 | 26,204,569 | 19.4% | 0.87% | 0% | 80.6% | 18.5% | 18.1% | 0.0353% | 0% | 0.0000305% |
| FDVT_2 | 19,625,150 | 18.8% | 0.86% | 0% | 81.2% | 17.9% | 17.4% | 0.0375% | 0% | 0.0000408% |
| FDVT_3 | 27,173,465 | 15.6% | 1.44% | 0% | 84.4% | 14.2% | 13.8% | 0.0412% | 0% | 0.0000626% |
| PSBB_1 | 30,555,168 | 23.9% | 0.165% | 0% | 76.1% | 23.7% | 23.5% | 0.0149% | 0% | 0.00000327% |
| PSBB_2 | 37,826,872 | 24.3% | 0.262% | 0% | 75.7% | 24% | 23.8% | 0.0178% | 0% | 0.00000529% |
| PSBB_3 | 26,924,648 | 23.5% | 0.166% | 0% | 76.5% | 23.3% | 23.1% | 0.015% | 0% | 0.0000111% |
| PSVT_1 | 26,247,423 | 19% | 0.421% | 0% | 81% | 18.6% | 18.4% | 0.0192% | 0% | 0.0000152% |

**Table S2.** Analysis of variance (ANOVA) of qPCR measurements.

| **a. Fixed-effects ANOVA using the 16S rRNA gene (value as the criterion)** |  |  |  |  |  |  |  |
| --- | --- | --- | --- | --- | --- | --- | --- |
| Predictor | Sum of squares | df | Mean Square | F | p | partial $\eta^2$ | partial $\eta^2$ 90% CI [LL, UL]<sup>a</sup> |
| (Intercept) | 245.55 | 1 | 245.55 | 1616.98 | .000 |  |  |
| 16S rRNA gene | 5.87 | 3 | 1.96 | 12.89 | .000 | .66 | [.35, .74] |
| Error | 3.04 | 20 | 0.15 |  |  |  |  |

| **b. Fixed-effects ANOVA using the *sul1* gene (value as the criterion)** |  |  |  |  |  |  |  |
| --- | --- | --- | --- | --- | --- | --- | --- |
| Predictor | Sum of squares | df | Mean Square | F | p | partial $\eta^2$ | partial $\eta^2$ 90% CI [LL, UL] |
| (Intercept) | 56.87 | 1 | 56.87 | 960.06 | .000 |  |  |
| *sul1* | 6.39 | 3 | 2.13 | 35.95 | .000 | .85 | [.68, .89] |
| Error | 1.13 | 19 | 0.06 |  |  |  |  |

| **c. Fixed-effects ANOVA using the *bla*<sub>CTX-M</sub> gene (value as the criterion)** |  |  |  |  |  |  |  |
| --- | --- | --- | --- | --- | --- | --- | --- |
| Predictor | Sum of Squares | df | Mean Square | F | p | partial $\eta^2$ | partial $\eta^2$ 90% CI [LL, UL] |
| (Intercept) | 0.00 | 1 | 0.00 | 0.00 | 1..0 |  |  |
| *bla*<sub>CTX-M</sub> | 5.70 | 3 | 1.90 | 17.09 | .000 | .72 | [.45, .79] |
| Error | 2.22 | 20 | 0.11 |  |  |  |  |

<sup>a</sup> LL and UL represent the lower-limit and upper-limit of the partial $\eta^2$ confidence interval, respectively.

